# Single-cell mass accumulation reveals bacterioplankton growth rate in native seawater

**DOI:** 10.1101/2025.08.08.669205

**Authors:** Yanqi Wu, Benjamin R.K. Roller, Cathrine Hellerschmied, Andreas Sichert, Annika L. Gomez, Nina Bartlau, Joana Séneca, Edo Danilyan, Michael Wolfram, Marc Mußmann, Teemu P. Miettinen, Martin F. Polz, Scott R. Manalis

**Affiliations:** Koch Institute for Integrative Cancer Research, Massachusetts Institute of Technology, Cambridge, United States; Centre for Microbiology and Environmental Systems Science, Division of Microbial Ecology, University of Vienna, Vienna, Austria; Institute of Molecular Systems Biology, ETH Zurich, Zurich, Switzerland; Department of Civil and Environmental Engineering, Massachusetts Institute of Technology, Cambridge, United States; Joint Microbiome Facility of the Medical University of Vienna and the University of Vienna, Vienna, Austria; Department of Biological Engineering, Massachusetts Institute of Technology, Cambridge, United States; Department of Mechanical Engineering, Massachusetts Institute of Technology, Cambridge, United States

## Abstract

The growth of marine microbial communities drives biogeochemical cycling of carbon and other elements, yet the growth rates of individual species within complex ocean ecosystems remain poorly understood. In particular, the coexistence of a large diversity of copiotrophic bacteria, which are capable of fast growth but typically remain at low abundance, has been interpreted as a feast or famine existence. Here we show that contrary to the notion of infrequent growth, *Vibrio* bacteria exhibited consistent growth rates in coastal ocean samples, despite representing only a small fraction of the total community. These observations were enabled by a suspended microchannel resonator (SMR), which we adapted to function as a single-cell chemostat. By maintaining a continuous supply of native seawater around each trapped cell, we prevented nutrient depletion and used the SMR’s high mass precision to resolve growth rates that are otherwise undetectable. *Vibrio* species displayed significantly larger cell mass and faster growth than other community members across samples collected at different temporal intervals from days to years. Surprisingly, their growth was consistently limited by carbon, contrary to the expectation that heterotrophic bacteria in the euphotic zone would be limited by nitrogen and phosphorus due to competition with algae. The correlation between cell mass and growth rate of *Vibrionaceae* in seawater followed established growth laws derived from laboratory conditions, suggesting that growth physiology observed in pure cultures is applicable to wild bacterial populations. Overall, our findings suggest that rare species may play a disproportionately large role in the marine carbon cycle, with rapid biomass turnover driven by a combination of high growth rates balanced by intense predation.

## Introduction

Microbes in the oceans employ different strategies to adapt to the prevailing conditions of dissolved nutrient scarcity, which are punctuated by nutrient hotspots of varying spatial and temporal dynamics. Oligotrophs —bacteria adapted to compete under extremely low nutrient conditions—dominate the global oceans in terms of absolute cell number, but their cells are small and metabolic activity is low, enabling them to eke out a living under severe nutrient limitation (*1*). In contrast, copiotrophs—bacteria that have the potential of rapid growth in nutrient-rich conditions—make up much of the genomic diversity in microbial communities and are thought to engage in feast-and-famine cycles, persisting in a starved state until encountering a short-lived nutrient pulse that fuels rapid growth (*1–3*). Because copiotrophs are typically rare in samples, it is often assumed that starvation is their predominant state. For example, species within the genus *Vibrio* are capable of some of the highest growth rates observed in pure cultures but are typically present at relative abundances of less than 1% of the total community (*4*). Recent studies have, however, shown that some copiotrophic taxa can contribute disproportionately to overall community respiration (*5*) and that growth can co-occur with little net change in abundance, presumably due to being offset by predation (*6*). While these results suggest a more important contribution of copiotrophs, it remains unknown if the observed activities represent “jackpot” events—where nutrient hotspots were sampled—or if rare copiotrophs can sustain high growth rates consistently and thereby contribute disproportionately to biogeochemical cycling due to their biomass turnover over longer time periods.

A major obstacle to understanding how specific bacterial populations contribute to biogeochemical cycles is the difficulty of directly measuring their growth in natural environments. Most current approaches rely on indirect proxies of biomass synthesis or metabolic activity. For example, community-wide estimates based on the incorporation of nucleotides or amino acids into biomass suggest that average bacterial division rates in the ocean range from 0.1 – 1 day⁻¹ (*2*). Although recent studies have provided a more differentiated picture with increased taxonomic resolution achieved using advanced fluorescence *in situ* hybridization (FISH) labeling (*6*), combined single cell respiration and genomics (*5*), or amplicon sequencing-derived techniques (*7*, *8*), these methods still face key limitations. Specifically, these approaches are biased towards more abundant groups and often need to average over broad taxonomic groups due to difficulties in sampling sufficient individuals or observing statistically significant changes in populations persisting at lower abundances. Similarly, dilution cultures that have been used to measure growth in seawater can only assess relatively abundant species and it is difficult to know potential limitations in nutrients that can arise when cell numbers start to increase. In light of these limitations, we reasoned that a promising path forward is to adapt a single cell technology—capable of directly measuring biomass accumulation—to the complexities of environmental samples. Biomass accumulation provides the most direct and interpretable measurement of microbial growth and it can be readily incorporated into biogeochemical models. We propose that copiotrophic bacteria, whose ecological roles remain poorly understood due to their typically low abundances, offer an ideal test case for applying such a single-cell approach.

In this study, we carry out the most direct measurement of growth by tracking mass increases of individual cells in their native seawater and identify the nutrients that limit this growth. To achieve this, we adapted the suspended microchannel resonator (SMR) (*9–11*), a microfluidic mass sensor capable of exceptionally precise single-cell mass measurements, for continuous monitoring of individual cells supplied with unamended, filtered seawater. By engineering the SMR system to maintain a constant flow of seawater around each trapped cell, we created a chemostat-like microenvironment that prevents nutrient depletion and reflects the ambient dissolved nutrient conditions at the time of sampling. This setup enables us to resolve mass accumulation and growth rates that are too subtle to detect with existing approaches, particularly in nutrient-poor environments. Because the filtered seawater lacks particles and other organisms, observed growth depends solely on the availability of dissolved organic and inorganic compounds. This platform thus allows for real-time, high-resolution tracking of bacterial growth under ecologically relevant conditions and independent of confounding factors that influence population-level growth such as viral or protozoan predation.

We began by measuring the growth rates of individual copiotrophic species in unamended seawater, both collected from the same coastal location. We found that two *Vibrio* species consistently exhibit faster growth rates than most other species. We then focused on *Vibrio cyclitrophicus* 1G07 as a model organism, as it represents a recently speciated, exclusively free-living lineage that does not attach to particles (*12*, *13*), making it particularly suitable for assessing growth based solely on dissolved nutrient uptake. We observed that this organism consistently grows at similar rates in seawater samples from different days and years and its variation in growth rate is correlated to single cell mass. We also found that the distribution of single-cell masses *in situ* can predict growth rates under natural conditions. Overall, our results challenge the prevailing “feast-and-famine” paradigm for copiotrophic bacteria.

## Results

### Direct measurement of single-cell mass accumulation using SMR

The SMR is a vibrating micro-cantilever containing an internal fluidic channel; when a cell passes through, its buoyant mass causes a measurable shift in the cantilever’s resonant frequency (**Fig. 1A**). By repeatedly flowing a single cell back and forth through the cantilever, we can track its mass over time and directly quantify growth. Buoyant mass—defined by the product of cell volume and the density difference between the cell and surrounding fluid—is a well-established proxy for dry mass, and can be converted to dry mass by a scaling factor (multiply by approximately 3) (*14–16*). For simplicity, we refer to buoyant mass as mass throughout. Growth rate is derived by first calculating the mass accumulation rate (the slope of mass vs. time) and then normalizing it by the cell’s mass to obtain the specific growth rate (**Fig. 1B**). The precision increases with trapping time, which we systematically characterized (**fig. S1** and **supplementary text**). For example, the mass accumulation of *Vibrio cyclitrophicus* 1G07 growing in nutrient-rich Marine Broth 2216 can be resolved with just 5 minutes of trapping (**Fig. 1C**). Importantly, we found that 30-minute trapping enables accurate growth measurements even for bacteria with doubling times on the order of weeks. We therefore used 5-minute trapping for fast-growing cells in culture media and 30-minute trapping for slow-growing cells in native seawater.

**Fig. 1.**
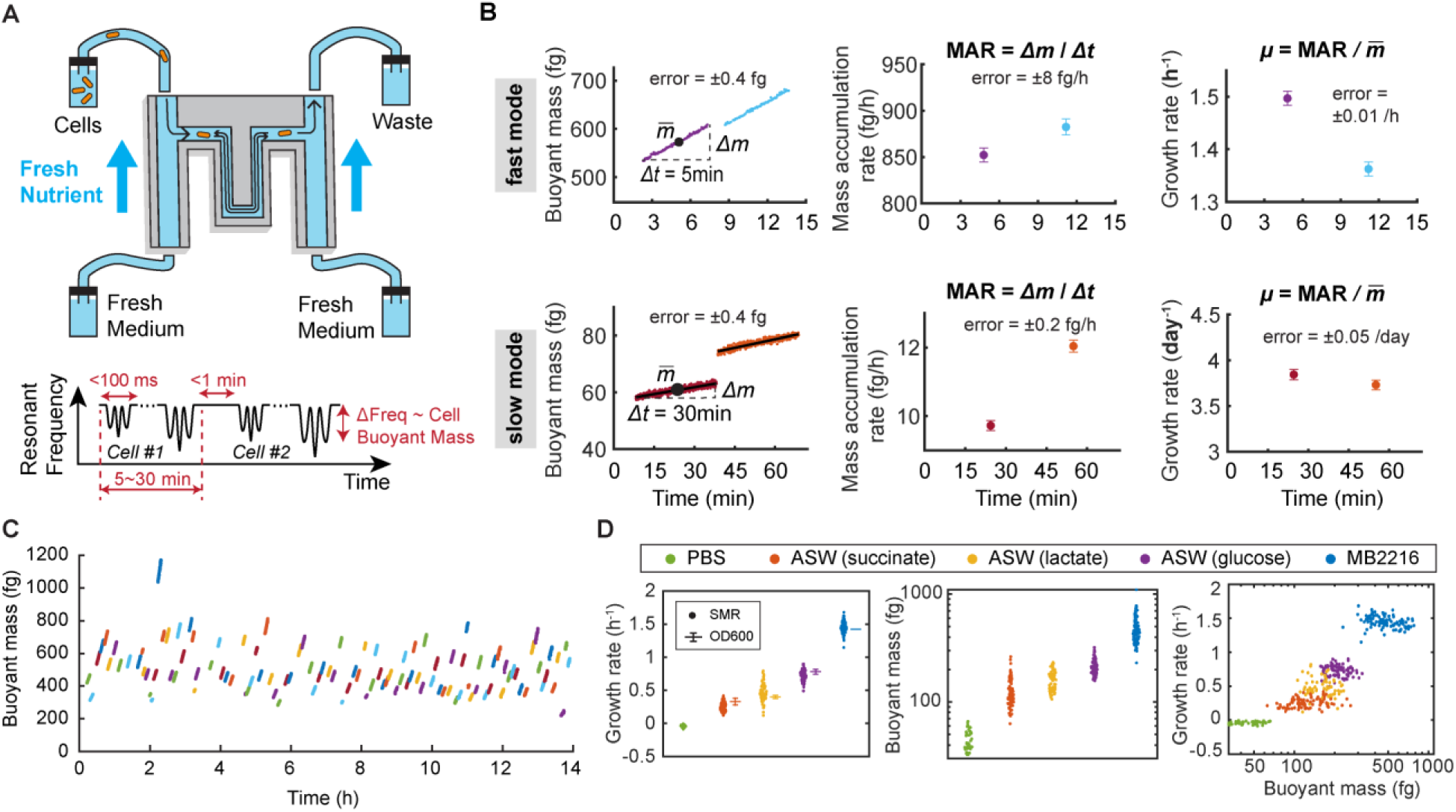
Suspended microchannel resonator (SMR) measures single-cell growth rate by continuous mass measurements. **(A)** SMR, comprising a vibrating cantilever with an internal microfluidic channel, can trap a single bacterium, by passing it through the cantilever repeatedly over a set time window while fresh nutrients are provided from the fresh medium vials. During the cell trapping, resonant frequency shifts of the SMR, caused by each traversal of the trapped cell, are captured in real time and converted to cellular buoyant mass. Once the current trapping measurement is completed, the system flushes the cell to the waste vial and searches for another cell from the cell vial within minutes. **(B)** Fast and slow modes that differ in trapping time (5 min vs 30 min). Fast mode is exemplified by *Vibrio cyclitrophicus* 1G07 in Marine Broth 2216, and slow mode by *Vibrio cyclitrophicus* 1G07 in filtered seawater. In both modes, continuous mass measurements are converted to an estimate of mass accumulation rate (MAR) and specific growth rate (*µ*). The measurements yield error levels of mass: 0.4 fg, MAR: 8 fg·h^-1^ (fast mode) or 0.2 fg·h^-1^ (slow mode), and specific growth rate *µ*: 0.01 h^-1^ = 0.24 day^-1^ (fast mode) and 0.05 day^-1^ (slow mode). **(C)** Representative single-cell growth curves of bacteria *V. cyclitrophicus* 1G07 in Marine Broth 2216. Each line represents 5-min trapping of a bacterium. **(D)** Mass and growth rate distributions of *V. cyclitrophicus* in different laboratory culture conditions, compared to population-mean growth rate measured by OD600 (mean ± standard deviation of replicate experiments). The conditions include phosphate buffered saline (PBS) as a negative control, artificial seawater (ASW) with succinate, lactate, or glucose as carbon source, and Marine Broth 2216 (MB2216).

We validated the performance of our SMR platform by comparing it to standard laboratory growth measures and evaluating potential biases. Growth rate estimates of *V. cyclitrophicus* 1G07 in different laboratory media growing at population-level rates from 0.3 to 1.5 h^-1^ were comparable between the SMR and OD600 measurements (*R^2^* = 0.98, **Fig. 1D**). Unlike bulk measurements, however, the SMR also provides paired mass and growth rate at the single-cell level, revealing additional insights. For example, we observed substantial heterogeneity in both cell mass and growth rate even in homogeneous media, with coefficients of variation averaging 24% and 19%, respectively. While population-average cell mass strongly correlated with growth rate across nutrient conditions (*R^2^* = 0.97), there was no detectable correlation between single-cell mass and growth rate within individual populations (*R^2^* < 0.05 in all cases; **Fig. 1D**). We further confirmed that the SMR’s continuous medium supply supports sustained growth, mimicking key features of a miniaturized chemostat (**fig. S2**), and that the system introduces minimal bias from 30-minute trapping or from chemical contamination (**fig. S3, S4** and **supplementary text**). Together, these results demonstrate that the SMR can accurately and robustly measure bacterial growth rates at single-cell resolution.

### Growth and nutrient limitation patterns of bacterial species in coastal seawater

To resolve the growth behaviors of diverse bacteria in their native environment, we spiked in individual species isolated from a coastal site (Nahant, MA, US East Coast) into filter-sterilized seawater collected from the same site in June 2022 and measured their growth in the SMR. Before the spike-in, these pre-grown bacteria were washed and diluted into the filtered seawater to avoid carryover of nutrients, as validated by our negative control experiments (**fig. S4** and **supplementary text**). After the spike-in, the bacteria acclimated to the seawater in the first several hours and then reached a steady state, where individual bacteria gained mass, and the population maintained stable mass and growth rate profiles consistent with the chemostat-like capability of the SMR (**Fig. 2A** and **fig. S5**). Steady-state mass and growth measurements under these conditions served as the basis for all subsequent analyses. All experiments in filtered seawater were conducted at 22°C, a temperature we validated as having minimal impact on growth relative to *in situ* seawater temperatures (**fig. S6**).

**Fig. 2.**
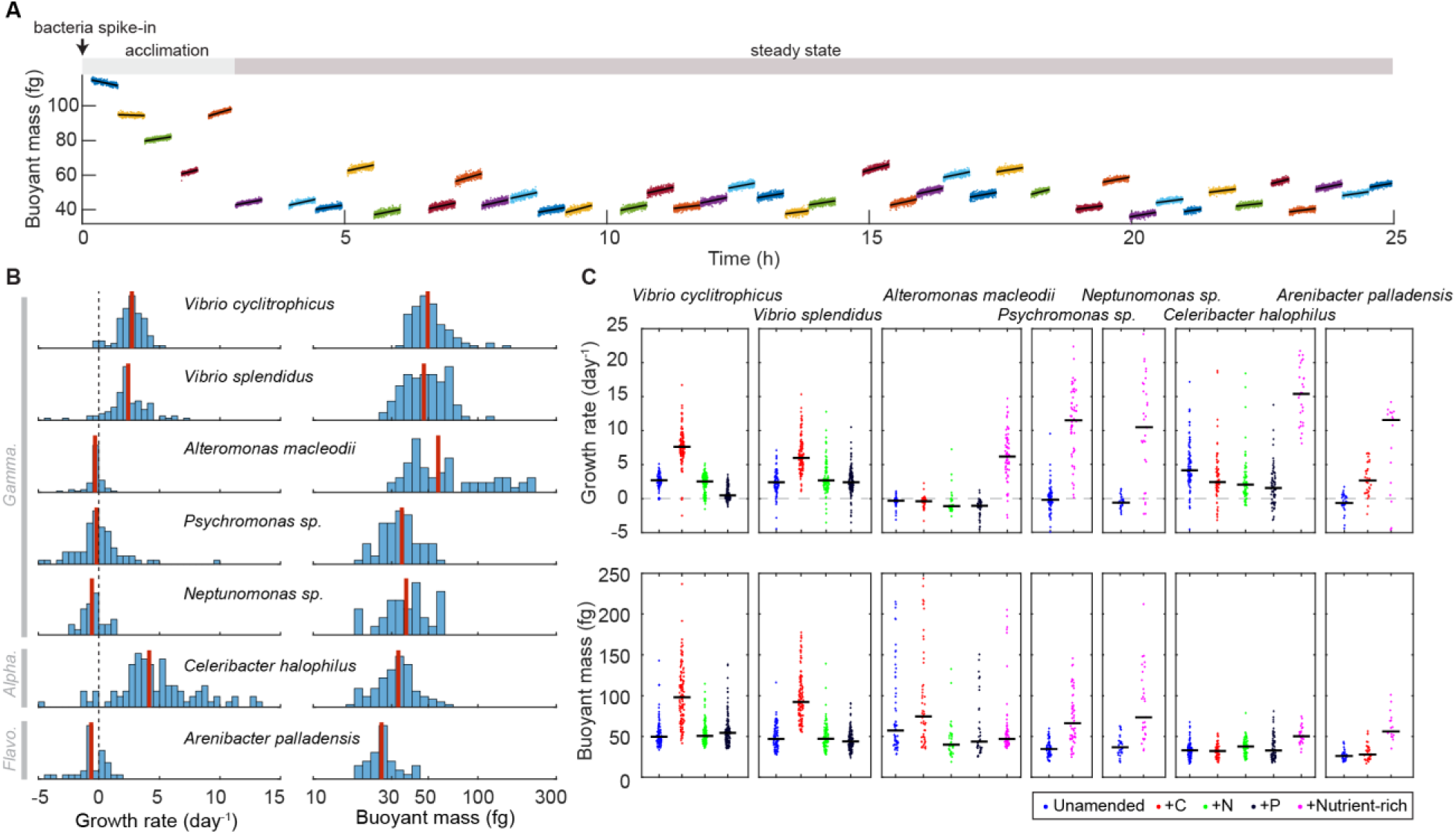
Growth rate and mass in coastal seawater are nutrient limited, but the response to additional nutrient availability differs across bacterial species. **(A)** Single-cell growth curves after bacteria were spiked into filtered seawater, reaching steady-state growth after an acclimation period, exemplified by *Vibrio cyclitrophicus* 1G07 in a coastal seawater sample on June 10, 2022. Each line indicates a 30-minute growth trajectory of individual bacteria. **(B)** Growth rate and mass distributions of different marine bacterial species in the seawater sample. All bacterial species were isolated from the same coastal location where the seawater sample was collected, and the class-level taxonomy was annotated as Gamma (*Gammaproteobacteria*), Alpha (*Alphaproteobacteria*) and Flavo (*Flavobacteriia*). The red vertical lines depict the median of the distributions and black vertical dotted line depicts zero growth rate for reference. **(C)** Growth rate and mass of different bacterial species in response to nutrient amendment of carbon (C = 20 µM glucose), nitrogen (N = 40 µM NH_4_Cl), phosphorus (P = 5 µM Na_2_HPO_4_) or chemically complex, nutrient-rich medium (“Nutrient-rich”). Each dot refers to a single cell datapoint and black lines refer to the median of the population.

We show significant variations in cell mass and growth across diverse bacterial species that were isolated from the same site and represent major phylogenetic groups in the ocean (including *Alphaproteobacteria*, *Gammaproteobacteria* and *Flavobacteriia* (*17*)). We found that several species (two *Vibrio* species and one *Celeribacter* species) grew at substantial population-average rates of 2.39–4.82 day^-1^, whereas the growth rates of the other species were indistinguishable from zero or negative, the latter possibly due to maintenance energy expenditure or death. The data thus suggest heterogenous growth and resource requirements across species (**Fig. 2B** and **table S1, S2**).

Although it is generally expected that in the euphotic zone bacteria should be nitrogen or phosphorus limited due to competition with algae (*18*, *19*), our results show that all species were either limited by organic carbon or some trace nutrients. We supplemented the same seawater sample with carbon (glucose), nitrogen (ammonium) or phosphorus (phosphate) at the maximum observed concentrations from public oceanic datasets (Hawaii Ocean Time-series (*20*, *21*), European seas (*22*, *23*), and Nahant Time-series (*24*)), or with chemically complex, rich nutrient mixture. Growth rates of all species significantly increased to 5.92–15.19 day^-1^ (*p*≤0.005) in response to either glucose or complex media but never in response to nitrogen or phosphorus (**Fig. 2C** and **table S1**). In the conditions of growth rate elevated by nutrient amendment, the cell mass also increased for all species (*p*≤0.002) except *Alteromonas macleodii*. In particular, both *Vibrio cyclitrophicus and V. splendidus* showed 3-fold growth rate increase and 2-fold cell mass increase in glucose amended seawater, but remained unchanged in N- and P-amended seawater or even decreased growth rate in one set of conditions. However, some species were not stimulated by glucose, N or P, but their growth increased when a mixture of nutrients was spiked, regardless of whether they exhibited growth in unamended seawater. This suggests that they required nutrient sources present in the complex mixture such as trace nutrients or organic carbon other than glucose.

### Consistent growth and carbon limitation of *Vibrio cyclitrophicus* over months and years

Because the relatively rapid growth of *Vibrio* in unamended seawater could be a result of an episodic growth spurt as expected under the feast-or-famine paradigm, we asked whether the growth and nutrient limitation patterns changed over longer time scales. Using *V. cyclitrophicus* 1G07 as a model, we measured single-cell growth rates in 10 seawater samples from 2010 and 2022 to ensure coverage of different conditions as they might occur over days to seasons and years (**Fig. 3A**).

**Fig. 3.**
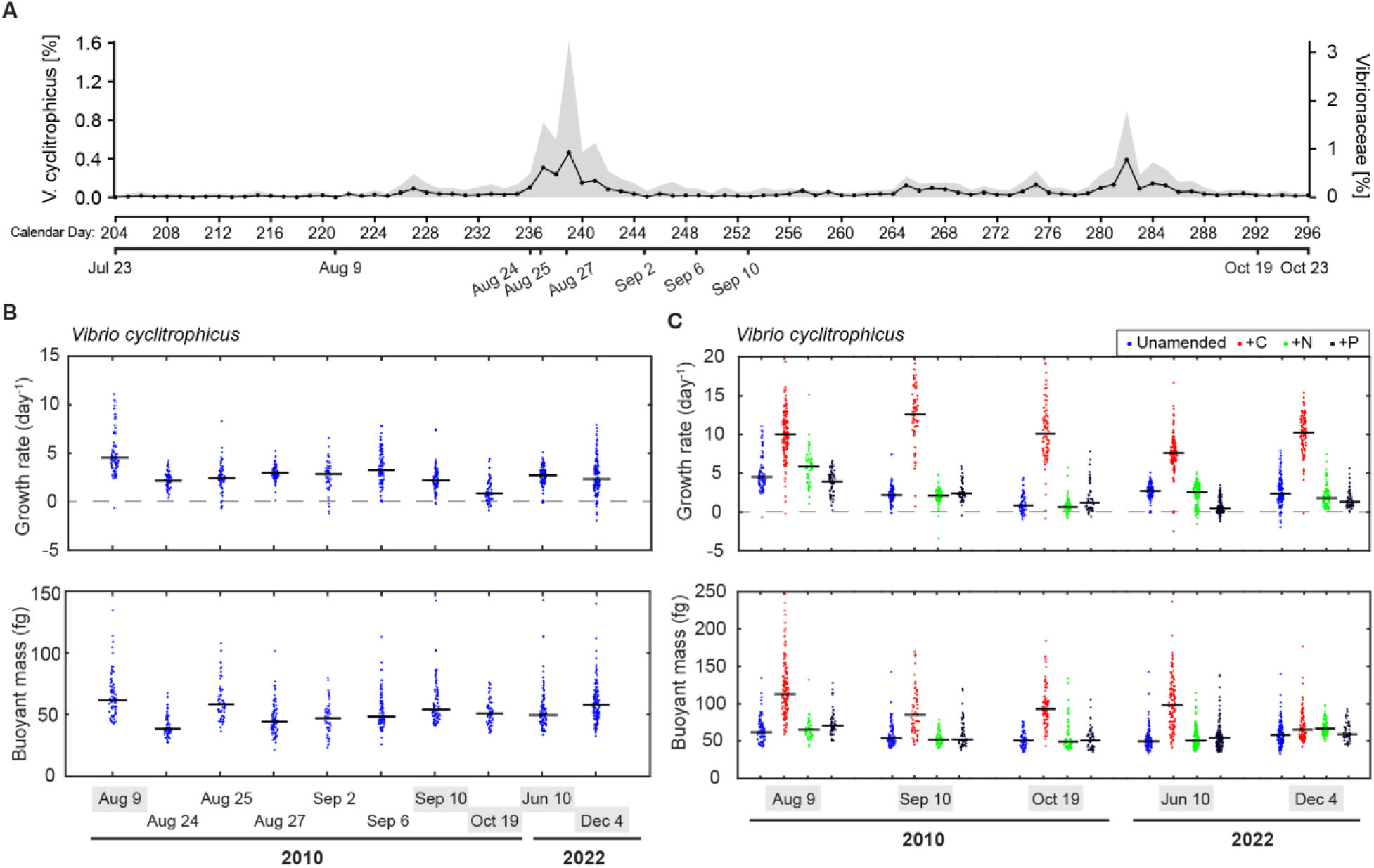
*Vibrio cyclitrophicus* exhibits consistent growth over months and years, even in conditions of low abundance. **(A)** Relative abundance of bacterial species *V. cyclitrophicus* (black dots and line) and family *Vibrionaceae* (gray shaded area) in the 93-day seawater samples collected in 2010. **(B)** Growth rate and mass of *V. cyclitrophicus* in ten representative unamended seawater samples in 2010 and 2022. The data for June 10 2022 has been shown in Fig. 2 and are repeated here. Five seawater samples were chosen and highlighted in gray for additional measurements in response to nutrient amendment shown in (C). **(C)** Growth rate and mass of *V. cyclitrophicus* in selected seawater samples with or without nutrient amendment of carbon (C = 20 µM glucose), nitrogen (N = 40 µM NH_4_Cl) or phosphorus (P = 5 µM Na_2_HPO_4_). Each dot refers to a single cell datapoint and black lines refer to the median of the population.

We found that *V. cyclitrophicus* grew in all unamended filtered seawater samples over these extended periods at approximately similar rates. The population-average growth rate mostly varied between 2 and 3 day^-1^ and the population-average cell mass varied around 50 fg (**Fig. 3B** and **table S1, S3**). The carbon amendments across seawater samples consistently increased the growth rate to 7.67–12.35 day^-1^ (*p*≤0.003 for each day) and the cell mass to 70.6–120.3 fg (*p*≤0.03 for each day, **Fig. 3C** and **table S1**). On the contrary, the nitrogen and phosphorus amendments did not increase the growth rate or cell mass (*p*=0.05– 0.97) and even decreased the growth rate or mass in some seawater samples. Importantly, because our time series includes near consecutive days without major decrease in growth rates and with the consistent pattern in carbon limitation, these observations suggest that carbon supply and consumption for these organisms are roughly balanced at this coastal site and that no major changes in growth physiology should occur from day to day.

To better understand which carbon sources support the growth of *V. cyclitrophicus* in filtered seawater, we measured the concentrations of 18 monosaccharides across three representative seawater samples (**table S4**). After 48 hours of incubation with *V. cyclitrophicus*, the concentrations of glucose and galactose decreased by 70% ± 9% and 35% ± 12% (n = 3), respectively, indicating selective uptake of these sugars (**fig. S7A**). To test whether glucose and galactose specifically limit growth, we supplemented a seawater sample with a mixture of seven monosaccharides, including these two. We again observed consumption of glucose and galactose, while concentrations of the other sugars remained largely unchanged—except for mannose, which unexpectedly increased (**fig. S7B, C and table S4**). This increase may result from oxidation of mannitol to mannose, which is not distinguishable using our current method. Together, these metabolomic data support the conclusion that the consistent growth of *V. cyclitrophicus* in seawater is fueled, at least in part, by the uptake of glucose and galactose.

Then we examined whether relative abundance was indicative of growth as generally believed in copiotrophs. We found that there was no obvious correlation between growth and relative abundance of *V. cyclitrophicus* in the samples from 2010, which have been fully metagenomically sequenced (*24*). For example, the fastest (∼5 day^-1^) and slowest growth (∼1 day^-1^) occurred on day 221 (Aug 9) and on day 292 (Oct 19) when the relative abundance of *V. cyclitrophicus* and *Vibrionaceae* family remained constantly low around 0.02% and 0.1% of the community. On the other hand, the growth rate was stable at 2–3 day^-1^ when both *V. cyclitrophicus* and the family experienced expansions around day 239 (Aug 27) reaching 0.5% and 3% of the community, respectively (**Fig. 3A, B**). Such decoupling of growth and relative abundance suggests that the dynamics of individual species are either masked by shifts of other species, or that top-down environmental forces, such as predation, regulate abundance. Overall, the results suggest that *V. cyclitrophicus* showed consistent rates in seawater despite their rarity in the community, contrary to the expectation of long-term starvation, and persistent carbon-limitation of their growth rates.

### Nutrient growth law links *in situ* cell mass to growth

We observed that the average mass of *Vibrio* cells increased in samples which supported faster growth rates across native and nutrient amended seawater. Such a relationship has been observed in controlled laboratory cultures for several model organisms and is commonly referred to as the nutrient growth law, which states that the mass of cells exponentially increases with growth rate (*25–29*). However, it is currently not clear if the slope of this relationship changes at slow growth rates (*28*). We therefore first sought to test whether *Vibrio* cells grown in native and amended seawater follow the same mass accumulation trends as those grown in richer laboratory media, and whether the growth law can be used to predict *in situ* growth rates based on mass distributions of cells sorted directly from seawater samples.

To establish a nutrient growth law generalizable for the broader *Vibrionaceae*, we selected 11 unique seawater isolates representing 5 *Vibrio* species grown in 9 different laboratory media (**Fig. 4A** and **table S5**). Treating each filtered seawater sample like an independent growth medium, we observed that the average mass and growth rate of *V. cyclitrophicus* growing in unamended or amended seawater follows a growth law with a slope that is not significantly different from the growth law measured only in laboratory culture media (**Fig. 4A**). The supplementation of seawater with carbon lead to cell sizes and growth rates that nearly perfectly follow the trend seen in laboratory media and are substantially faster than the slowest growth rates achieved in laboratory media. From these results we conclude that a single nutrient growth law applies across all growth rates observed in seawater and in the laboratory.

**Fig. 4.**
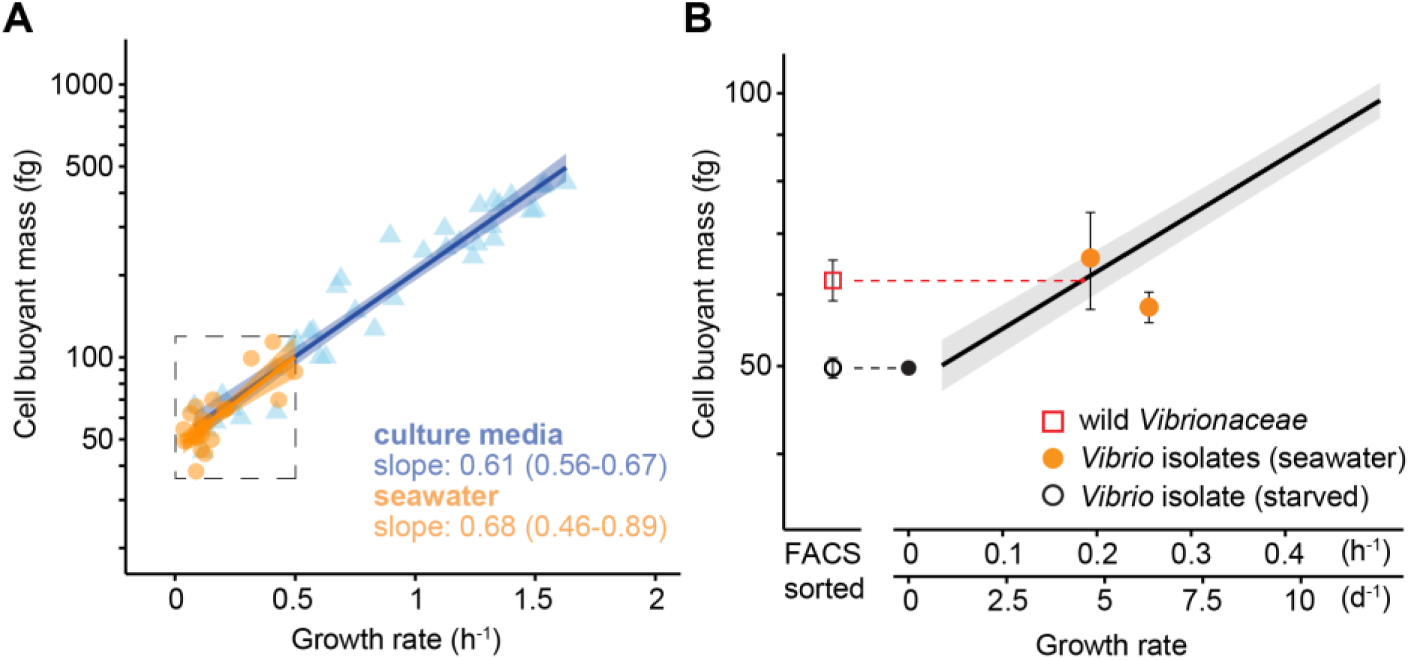
A nutrient growth law governs the average mass and growth rate of *Vibrionaceae* growing in laboratory media and in seawater. **(A)** The average mass and growth rate of 11 *Vibrionaceae* strains representing 5 species and grown in 9 different media (blue triangles, n = 38 independent strain/media combinations) has a slope that is statistically indistinguishable from *Vibrio cyclitrophicus* grown in seawater from 10 different days with or without nutrient amendment (orange circles, n = 25 independent day/amendment combinations). 95% confidence intervals reported in parentheses overlap when separate regression models for seawater and culture media (each with a single covariate of growth rate). When a multiple regression model is fit for a combined seawater and culture media dataset (with sample-type and its interaction with growth rate included as covariates) there is no evidence for either term being significant (sample-type: *p* = 0.349; interaction: *p* = 0.583). **(B)** The *in situ* mass of wild *Vibrionaceae* cells sorted from complex seawater communities (red square, n = 3 replicate seawater samples, 1175 total cells) corresponds to the nutrient growth law (black line, regression of all seawater and laboratory media data points in panel (A). The cell mass and growth rates of two *Vibrionaceae* isolates grown in paired filtered seawater from the same sample in the SMR (orange dots) are included for reference and are used to approximate plausible limits for the *in situ* growth rate estimate. Starved *V. cyclitrophicus* cell mass is included (open black circle, n = 3 replicate cultures) and mapped to the nutrient growth law at zero growth rate for reference (filled black circle). All error bars on data points represent 1 standard deviation above and below the average mass.

To test whether wild *Vibrionaceae* populations reflect the masses and growth rates of cultivated cells in filtered seawater, we collected seawater from the same coastal site in 2023 and prepared two types of samples: filtered seawater (0.2 µm) for growth assays, and fixed cells from the 0.2–1 µm biomass fraction for direct analysis. We used fluorescence *in situ* hybridization (FISH) to label *Vibrionaceae* cells, followed by fluorescence-activated cell sorting (FACS) to isolate them from the broader bacterial community (**fig. S8**). Sorting accuracy was verified by amplicon sequencing (**fig. S9**) and sample fixation with ethanol resulted in an estimated 18% mass loss (**fig. S10** and **supplementary text**). The corrected median mass of sorted wild *Vibrionaceae* cells was 62.1 fg which was substantially higher than that of a starved, non-growing *Vibrio* control subjected to the same fixation, labeling, and sorting protocol (48.5 fg). This measured cell mass for wild *Vibrionaceae* cells corresponds to an inferred growth rate of 4.39 day^-1^ based on the growth law equation, aligning closely with growth rates (4.64 & 6.12 day^-1^) observed in two *Vibrio* species cultured in the paired filtered seawater in the SMR (**Fig. 4B**). Although bacterial populations in the wild experience a more heterogeneous environment than filtered seawater, experiencing small scale nutrient patches, competition, and predation, our results nonetheless suggest that the growth rates measured in filtered seawater approximate the actual growth of planktonic *Vibrionaceae* in the wild.

## Discussion

The SMR has enabled precise, direct measurements of bacterial growth rate at the single-cell level in native seawater, revealing physiological dynamics that are difficult or impossible to access with existing methods. We observed that copiotrophic taxa, such as *Vibrio*, consistently sustain growth rates equal to or even faster than the global coastal community average and commonly studied oligotrophs like the SAR11 clade (**fig. S11**). For these fast-growing bacteria to remain at low relative abundance, they must be subject to strong top-down regulatory pressures such as predation, potentially due to traits that make them more “edible” or energetically favorable targets (*30*). The coexistence of rapid growth and low abundance suggests that *Vibrio* and perhaps other copiotrophs have a disproportionate influence on ecosystem function, particularly in carbon cycling, by rapidly turning over biomass even when numerically rare. These insights not only challenge assumptions about the ecological roles of rare taxa but also offer a path toward refining ecosystem models by integrating microbial identity, physiology, and ecological impact.

The bacterial growth law has been proposed to describe the relationship between cell mass and growth rate under balanced, steady-state growth (*25*, *31*). It is often assumed that physiology in natural environments is far from steady state, because microbial growth is shaped by fluctuating conditions and resource limitations. Despite this common assumption, *Vibrio* populations grew at similar rates across near-consecutive days and followed a nutrient growth law indistinguishable from that observed in laboratory media. This suggests that, at least in this coastal system, the balance between nutrient supply and consumption for planktonic cells may be more stable than previously appreciated. Consequently, physiological principles derived from controlled experiments—including cell mass as a proxy for growth rate—may hold predictive value in the wild. Moreover, our measurements of wild *Vibrionaceae* mass (**Fig. 4B**) were performed using a commercially available SMR platform, suggesting that similar approaches could be adopted more broadly. When combined with molecular data, such mass-based physiological measurements may provide a more complete and scalable understanding of microbial life and activity in natural ecosystems.

## Materials and Methods

### SMR setup, operation and measurement resolution

The suspended microchannel resonator (SMR) consists of a vibrating micro-cantilever with an embedded fluidic channel that enables flow-through mass measurements of single bacterial cells. The fabrication and setup of the SMR followed previously established protocols (*9*, *32*). In this study, we used two types of SMR systems: a commercially available platform (LifeScale, Affinity Biosensors) and a custom-built setup designed for single-cell growth measurements (*10*). The LifeScale performs high-throughput buoyant mass measurements by automatically drawing samples from a well plate and passing them through the SMR sensor, enabling flow-through analysis of thousands of cells per minute. In contrast, the lab-built system enables repeated measurements of individual cells. Using computer-controlled pressure, the same cell is alternately pushed back and forth through the cantilever, allowing sub-100 millisecond mass measurements interspersed with 2–10 second holding periods. Each cell is “trapped” in this manner for 5–30 minutes before being flushed and replaced. Continuous perfusion of fresh medium from both ends of the cantilever maintains a stable nutrient environment throughout the trapping period. Each experiment lasted up to 24 hours and typically included 20–100 cells. From these measurements, we calculated cell mass (average mass over the trapping period), mass accumulation rate (slope of mass vs. time), and specific growth rate (mass-normalized accumulation rate). Additional details on resolution, validation, and control experiments are provided in the supplementary text.

### Seawater sampling

All coastal seawater samples were collected at Canoe Beach, Nahant, MA, USA (Lat: 42° 25′ 10.6″ N, Lon: 70° 54′ 24.2″ W). The 2010 sampling was detailed in a previous report (*24*), and the 2022 & 2023 sampling followed similar approaches with three replicates spatially separated along the bay. For 2010 sampling, 4 L of seawater were filtered through a 0.2 µm Sterivex filter membranes. The membranes were stored at −20°C until nucleic acids were extracted using a bead beating approach for further analysis. For 2010, 2022, and 2023 sampling, additional seawater was filtered to remove all microorganisms (0.7 µm GF/F for 2010, 0.2 µm polycarbonate for 2022, and 0.2 µm polyethersulfone for 2023) and the filtered seawater was stored at −80 °C prior to use in SMR growth experiments. In 2023, 1L of seawater biomass was sequentially fractionated, first through a 63 µm plankton net and then using vacuum filtration through 47 mm polycarbonate filter membranes of decreasing pore size (5 µm, 1 µm, and 0.2 µm, Whatman Nuclepore). Filter membranes containing fractionated biomass were flash frozen on dry ice and stored at −80°C. In 2010 & 2023 sampling, air and water temperatures were measured by a thermometer at the time of sampling, whereas seawater temperature in 2022 was obtained from long-term datasets of NOAA Station 44013, located 16 nautical miles east of Boston, MA and representing the regional conditions. Bacterial isolates were collected at the same site, minimally passaged, and kept at −80 °C. Details for bacterial isolation were previously reported (*33*, *34*).

### Laboratory culture media

To achieve growth rates between 1.6 h^-1^ and 0.05 h^-1^, 9 types of media were employed: Marine broth 2216 (MB2216), artificial seawater (ASW) + 10 mM glucose + amino acids (EZ) + nucleotides (ACGU), ASW + 10 mM glucose + EZ, ASW + 10 mM glucose, ASW + 20 mM L-lactate + EZ, ASW + 20 mM L-lactate, ASW + 20 mM L-succinate, ASW + 40 mM L-alanine, and ASW + 20 mM L-glutamate. MB2216 (Difco, 1x) was dissolved in ultra-pure water and autoclaved in a standard autoclaving cycle at 121 °C for 15 minutes. Autoclaved MB2216 medium was sterile filtered using a 0.1 µm filtration system (Sartorius Minisart Filter Unit, Millipore® Stericup® Quick Release Vacuum Filtration System). All ASW media were prepared from HEPES buffered artificial seawater base to which nitrogen (9.3 mM ammonium chloride), phosphorus (0.28 mM Di-sodium hydrogen phosphate), vitamins, trace elements and a varying carbon source were added to modulate growth rate. D-Glucose, L-Lactate, L-Succinate, L-Alanine, L-Glutamate (pH 8) were prepared as 1M, filter sterilized stock solutions (0.1 µm) and each diluted into the ASW medium to a specific concentration as stated above. To achieve growth rates in the intermediate to fast regime, commercially available solutions of 1x amino acids (EZ, Teknova, Holister, CA, USA) or 1x nucleotides (ACGU, Teknova, Holister, CA, USA) was supplied to the ASW medium. The ASW base was prepared in ultra-pure water and contains molar concentrations of the following anhydrous and hydrous salts: 410 mM NaCl, 280 mM Na_2_SO4, 9 mM KCl, 0.84 mM KBr, 0.4 mM H_3_BO_3_, 4.7 × 10^-2^ mM NaF, 10 mM MgCl_2_·6H_2_O, 10 mM CaCl_2_·2H_2_O, 9 × 10^-2^ mM SrCl_2_·6H_2_O, 24 mM NaHCO_3_ and 54 mM HEPES. The pH of the ASW base was adjusted to 8.1 (± 0.05) with 5 M sodium hydroxide and sterilized by autoclaving followed by filtration through a 0.22 µm membrane (Sartoclear Dynamics® Lab V 1000 mL Kit, 0.22 µm pore Size) to remove precipitates.

The vitamin and trace element solution were prepared as a 1000 X stock solution and contained 50 mg /L 4-aminobenzoic acid, 100 mg/L pyridoxine-HCl, 50 mg/L Thiamine-HCl, 50 mg/L Riboflavin, 50 mg/L Nicotinic acid, 50 mg/L di-Ca pantothenate, 50 mg/L Lipoic acid, 50 mg/L Nicotinamide Acid, 50 mg/L Vitamin B12, 20 mg/L Biotin, 20 mg/L Folic acid and 4 g/L FeCl3 dissolved in 6.5 mL 32% HCl, 2 g/L EDTA, 70 mg/L ZnCl_2_, 100 mg/L MnCl_2_·4H_2_O, 120 mg/L CoCl_2_·6H_2_O, 120 mg/L CoCl_2_·6H_2_O, 15 mg/L CuCl_2_·2H_2_O, 25 mg/L NiCl_2_·6H_2_O and 25 mg/L Na_2_MoO_4_·2H_2_O respectively. Both vitamin and trace element stock solutions were stored in dark, light tight containers at 4° C or −20 °C.

To improve stability of the ASW with respect to precipitation the HEPES salt was replaced by a 1M HEPES pH 8.0 sterile stock solution (Thermo Fischer Scientific) and autoclaving was omitted for some experiments. To ensure minimal background particle load, all experiments were performed in freshly prepared 0.1 µm PES filtered ASW media no older than 24 h. Particles below the limit of detection was removed on the LifeScale, but more particles could be formed spontaneously in the medium over the time span of the experimental procedures up to ∼ 10^5^ ml^-1^. Controls were always measured to verify background particles were less than 1% of measured cells.

### Batch growth experiments using *Vibrionaceae* isolates in laboratory culture media

To achieve the greatest reproducibility, all growth experiments of *Vibrionaceae* strains were performed at steady-state growth, determined by stable, exponential growth in OD600 and our previous recommendations for achieving steady-state mass in pure culture (*16*). *Vibrionaceae* strains (see table S5 for the strain list) were recovered from a freezer stock into MB2216 (Difco) and grown to stationary phase, after which they were diluted at least 10,000-fold into fresh laboratory culture medium of interest, through multiple 10-fold or 100-fold serial dilutions into the same fresh medium in sterilized glass tubes and incubated at 200 rpm and 25 °C. OD600 was monitored (Genesys 40, Thermo Scientific) and recorded approximately from 0.005 to 0.15. The cultures were sampled and fixed in steady state growth between OD600 0.05 and 0.15 for 1 h on ice with 4 % formaldehyde (Carl Roth, Lot no. 40334295). Fixed cells were stored in the native medium with the fixative at 4 °C until further use.

All cell mass measurements were performed in 1x PBS in ultra-pure water, prepared from 10x PBS stock solution (Fisher Scientific, Austria) and filtered across a 0.1 µm PES syringe filter membrane (Sartorius, Minisart Filter Unit). For each sample an aliquot of fixed cells was centrifuged at >16 000 g for 10 minutes, washed, and resuspended in 1x PBS. The samples along with controls of filtered medium were transferred to a 96-well plate (Deepwell Plate, Eppendorf) and analyzed on the LifeScale system for 5 minutes per sample.

### Fluorescence *in situ* hybridization, fluorescence activated cell sorting, and mass measurement of wild *Vibrionaceae*

Fluorescence *in situ* hybridization (FISH) and fluorescence activated cell sorting (FACS) were used to label and purify cells from wild *Vibrionaceae* populations for mass measurement, but these techniques could also impact the cell mass. In order to quantify the impact of FISH and FACS on cell mass, *V. cyclitrophicus* 1G07 was starved in 1x PBS for 20 hours. Since we knew their living mass, it provided a useful comparison for wild cells with unknown growth rates. The sampling and processing procedures used for fractionated biomass were applied to starved *V. cyclitrophicus* 1G07 samples. 3 replicates of starved *V. cyclitrophicus* 1G07 at 1×10^6^-1×10^7^ cells per ml were analyzed on the LifeScale (live sample) and concurrently filtered onto 47 mm polycarbonate 0.2 µm pore size filter membranes (Whatman Nuclepore), flash frozen on dry ice, and stored at −80 °C. Both starved *V. cyclitrophicus* 1G07 and fractionated seawater biomass samples (ranging from 0.2 µm - 1 µm due to previous filtration through 1 µm filter) on 0.2 µm polycarbonate filters were fixed with Ethanol. Filters were thawed and 1 mL molecular grade absolute ethanol pipetted directly to filter sitting on a vacuum filtration frit for 15 minutes. Vacuum was applied to remove ethanol and dried filters were cut using a sterile razor blade into technical replicate filter sections and stored at −80 °C.

FISH was performed on both starved cells and fractionated seawater biomass using a standard protocol for filtered samples (*35*) with some noted modifications. Filter pieces were thawed and placed on glass microscope slides using sterile forceps, then 30 µL of hybridization buffer containing FISH probes was applied directly on filter (either control mix or experimental mix), with probes each at 0.166 µM final concentration. Hybridization buffer contained 30 % formamide, as well as 0.1 g/mL dextran sulfate and 1 % blocking reagent to minimize probe binding to the filter. The sample was then incubated for 90 minutes at 46 °C inside a sealed conical tube with a paper towel wetted with excess hybridization buffer to control humidity during hybridization. After hybridization, filters were transferred into pre-warmed wash buffer at 48 °C using sterile forceps and incubated for 15 minutes. Washed filters were then briefly dipped in ice-cold ultrapure water for 3 seconds, then transferred into 1 mL of 150 mM NaCl + 0.05 % Tween80 resuspension buffer in low-bind microcentrifuge tubes previously shown to perform well for resuspending marine bacteria from filters (*36*). Tubes containing filters in resuspension buffer were incubated for 30 minutes at 37 °C, then vortexed horizontally at room temperature for 15 minutes at maximum speed, and filters removed from the resuspended cell solution, dried and stored in microcentrifuge tubes at −20 °C. Control probe mix contained two nonsense probes that should not bind any cells: NonEub338-1 (*37*) terminally linked to 2 fluorescein fluorophores per oligonucleotide and NonEub338-1 (*37*) terminally linked to 2 cyanine-5 fluorophores. Experimental probe mix contained two probes that should bind either bacterial cells or Vibrionaceae cells: Eub338-1 (*38*) (targeting most bacteria) linked to 4 fluorescein fluorophores per oligonucleotide and GV (*39*) (targeting *Vibrionaceae)* terminally linked to 2 cyanine-5 fluorophores. All probes purchased from biomers.net GmbH (Germany). The FISH probe sequences are: GV AGG CCA CAA CCT CCA AGT AG

Eub338-1 GCT GCC TCC CGT AGG AGT

NonEub338-1 ACT CCT ACG GGA GGC AGC

The mass of resuspended cells was measured on the LifeScale (presorting sample) and then cells were analyzed with FACS (BD FACS Melody) to sort putative Bacteria and *Vibrionaceae* populations based on two fluorescent phenotypes with experimental probes (Bacteria: fluorescein positive & cyanine-5 negative; Vibrionaceae: Fluorescein positive & cyanine-5 positive) or to putative false positive and true negative populations based on two fluorescent phenotypes with control probes (true negative: Fluorescein negative & cyanine-5 negative; false positive: fluorescein positive & cyanine-5 positive). Samples of *V. cyclitrophicus* 1G07 in pure culture that were either FISH stained or unstained were used to establish threshold settings on the fluorescein channel. These pure culture samples were also used to establish gates for sorting subpopulations using first the fluorescein vs. forward scatter plot & then the cyanine-5 vs. forward scatter plot to distinguish cells with a fluorescence signal (positive) from those without a fluorescence signal (negative). For seawater & pure culture samples labeled with either experimental probes or control probes, cells which were fluorescein positive were then further gated on cyanine-5 and sorted into cyanine-5 positive (experimental probes: *Vibrionaceae*; control probes: false positive) or cyanine-5 negative populations (experimental probes: Bacteria). For seawater and pure culture samples labeled with control probes, cells which were fluorescein negative were further gated into a cyanine-5 negative population (control probes: true negative). Gates and thresholds were adjusted so some signal was always detected for unstained samples or cell-free resuspension buffer in the fluorescein negative & cyanine-5 negative gates. Sorting fluid controls were collected by running a sample of cell-free resuspension buffer through the FACS machine and sorting the double negative gate to capture any unlabeled cells in either the sheath fluid or tubing that could potentially contaminate our sorted samples.

Sorted cells from all putative populations and controls were then split into three aliquots for either: 1) DNA extraction and sequencing, 2) imaging under 100x magnification on a Leica DM4B fluorescence microscope with Leica Thunder post-acquistion image processing, or 3) mass measurement on the LifeScale SMR (sorted sample). Sequencing and imaging were used to verify the cell sorting strategy worked as intended. DNA extraction of sorted cells, sorted controls, and presort samples was performed using the QIAamp DNA mini kit (Qiagen) following the manufacturer’s protocol with the exception of an additional enzymatic lysis step performed prior to the prescribed protocol. For this step, pelleted cells were resuspended in 180 µL of enzymatic lysis buffer (20mM Tris-chloride pH 8.0, 2mM Sodium EDTA, 1.2 % Triton X-100, 20 mg/ml lysozyme) and incubated for 30 minutes at 37 °C. DNA extraction of FISH stained filters (containing remnant biomass after cell resuspension) was performed using the DNeasy PowerSoil Pro kit. Extraction controls were performed with all reagents and kits, but excluding any input DNA. Sequencing was performed by the Joint Microbiome Facility of the University of Vienna and the Medical University of Vienna (project ID JMF-2505-03) using primers 341F/785R (*40*) and a unique dual-barcoding two-step PCR approach as described previously (*41*). Amplicon pools were sequenced on an Illumina MiSeq (2x 300bp). Raw data processing was performed as described previously (https://pubmed.ncbi.nlm.nih.gov/34093488/). Briefly, demultiplexing was performed with the python package demultiplex (Laros JFJ, github.com/jfjlaros/demultiplex) allowing one mismatch for barcodes and two mismatches for linkers and primers. Amplicon sequence variants (ASVs) were inferred using the DADA2 R package v1.42 (https://www.ncbi.nlm.nih.gov/pubmed/27214047) applying the recommended workflow (https://f1000research.com/articles/5-1492). FASTQ reads 1 and 2 were trimmed at 230 nt with allowed expected errors of 2. ASV sequences were subsequently classified using DADA2 and the SILVA database SSU Ref NR 99 release 138.2 (https://zenodo.org/records/14169026). The 16S rRNA gene amplicon sequencing data has been deposited at the Sequence Read Archive under the BioProject accession PRJNA1291099.

### Growth experiments using bacterial isolates in filtered seawater

Bacterial isolates were inoculated from freezer stock into ASW + 20 mM glucose. The cultures were shaken at 250 rpm and kept at room temperature (consistently 22 °C) unless specified otherwise. The cell density was continuously monitored by an OD600 photometer (MicrobeMeter, Humane Technologies). Once the cultures remained in stationary phase (OD of ∼1.0) for more than 2 hours, an aliquot was taken, washed twice in filtered seawater, and diluted 1:1,000 into filtered seawater either unamended or amended with additional nutrient to test for growth limitation. Amended seawater was supplemented with carbon (20 µM glucose), nitrogen (40 µM NH_4_Cl), phosphorus (5 µM Na_2_HPO_4_), or a chemically complex nutrient mixture (0.01x Marine Broth 2216 or artificial seawater + 10 mM glucose).

For growth measurement by SMR, the diluted culture in filtered seawater was immediately loaded into one vial of the SMR system and filtered seawater from the same sample and treatment was loaded into the other vials, allowing for measurements of cell mass and growth rate under continuous seawater supply. For characterizing monosaccharides in seawater, the diluted culture in filtered seawater was kept in a culture tube shaken at 250 rpm and aliquots were taken at 0, 0.5, 2, 4, 8, 12, 24, and 48 hours. As a negative control, aliquots were also taken from the same filtered seawater without any cells at 0, 2, 12, 24, and 48 hours. Each aliquot was centrifugated at 13,000 g for 5 minutes and the supernatant was taken and stored at −80 °C for further LC-MS analysis. The tested seawater were natural samples collected on Jun-10-2022, Aug-24-2010, and Oct-19-2010. One additional sample was prepared by supplementing the Jun-10-2022 seawater sample with fucose, galactose, glucosamine, glucose, mannose, rhamnose, xylose (10 nM each).

### Abundance of *Vibrio cyclitrophicus* and *Vibrionaceae* in time series metagenomes

The DNA of unfractionated samples was cleaned and size-selected using SPRI beads (AgenCourt® AMPure® XP, Beckman Coulter), libraries were prepared using a modified Nextera Flex kit and sequenced by the BioMicroCenter (MIT, Cambridge, MA) using an Illumina NovaSeq S1flowcell (2×150bp reads). A subset of these samples (days 222, 246, 252, 257, 259, 263, and 267) did not yield sufficient sequencing depth and were re-sequenced in 2022 at the Joint Microbiome Facility (University of Vienna, Vienna, Austria). Libraries of samples were prepared using the NEBNext® Ultra™ II FS DNA Library Prep kit (New England Biolabs) and sequenced on an Illumina NovaSeq SP flow cell (2×150 bp reads).

To determine the relative abundance of different taxonomic units in the metagenomes, the reads were processed as follows: (i) phiX sequences were removed with Bowtie2 (v. 2.3.4.2) (*42*) and quality trimmed with TrimGalore (v.0.5.0) (https://github.com/FelixKrueger/TrimGalore) with the parameters “-e 0.05 -- clip_R1 1 --clip_R2 1 --three_prime_clip_R1 1 --three_prime_clip_R2 1 --length 70 --stringency 1 –paired --max_n 1 --phred33”; (ii) error correction was done with Tadpole (v. 38.18, BBTools package https://jgi.doe.gov/data-and-tools/software-tools/bbtools/); (iii) reads were filtered using Kraken2 (v.2.1.3) (*43*) classification to retain only bacterial and archaeal sequences. In short, the Kraken database was built with reference libraries of archaea and bacteria, Kraken2 was run and reads were filtered with extract_kraken_reads.py (KrakenTools, https://github.com/jenniferlu717/KrakenTools); iv) Mapping to genomes was performed with bbmap (v.39.06) (BBTools package) with the parameters “fast=t minid=0.95 idfilter=0.98” and results sorted with samtools (*44*).

The relative abundances of *V. cyclitrophicus* were determined by competitively mapping metagenome reads from each day to all isolates identified as *V. cyclitrophicus* from the Nahant time series (**table S7**) and the RefSeq genome of *V. cyclitrophicus* (GCF_005144905.1). Mapped reads were then summed, and relative abundance calculated. Similarly, to determine the relative abundance of *Vibrionaceae*, the metagenome reads were competitively mapped to all available RefSeq genomes of the *Vibrionaceae* (218, NCBI, Nov. 2023) and summed. Plots were done with RStudio.

### Liquid chromatography-mass spectrometry (LC-MS) analysis of dissolved monosaccharides

Following a previously published protocol (*45*), the supernatant aliquots collected from bacterial culture in filtered seawater (25 µL) were derivatized with 75 µL of 0.1 M 1-phenyl-3-methyl-5-pyrazolone (PMP) in 2:1 methanol:ddH2O with 0.4 % ammonium hydroxide for 100 minutes at 70 °C. For quantification, we derivatized a serial dilution of a standard mix containing Galacturonic acid, D-Glucuronic acid, Mannuronic Acid, Guluronic Acid, Xylose, Arabinose, D-Glucosamine, Fucose, Glucose, Galactose, Mannose, N-Acetyl-D-glucosamine, N-Acetyl-D-galactosamine, N-Acetyl-D-mannosamine, Ribose, Rhamnose and D-galactosamine. Additionally, each sample and standard were spiked with an internal standard of 10 µM ^13^C_6_-Glucose, ^13^C_6_-Galactose and ^13^C_6_-Mannose (mass 186.11 Da). After derivatization, samples were neutralized with HCl followed by chloroform extraction to remove underivatized PMP as described previously (*46*).

Following Xu et al. (*46*), PMP-derivatives were measured on a SCIEX qTRAP5500 and an Agilent 1290 Infinity II LC system with a Waters CORTECS UPLC C18 Column, 90 Å, 1.6 µm, 2.1 mm X 50 mm reversed phase column with guard column. The mobile phase consisted of buffer A (10 mM NH_4_Formate in ddH2O, 0.1 % formic acid) and buffer B (100 % acetonitrile, 0.1 % formic acid). PMP-derivatives were separated with an initial isocratic flow of 15 % Buffer B for 2 minutes, followed by a gradient from 15 % to 20 % Buffer B over 5 minutes at a constant flow rate of 0.5 mL/min and a column temperature of 50 °C. The ESI source settings were 625 °C, with curtain gas set to 30 (arbitrary units), collision gas to medium, ion spray voltage 5500 (arbitrary units), temperature to 625 °C, Ion source Gas 1 &2 to 90 (arbitrary units). PMP-derivatives were measured by multiple reaction monitoring (MRM) in positive mode with previously optimized transitions and collision energies (*46*). Different PMP-derivatives were identified by their mass and retention in comparison to known standards. Technical variation in sample processing were normalized by the amount of internal standard in each sample. Peak areas of the 175 Da fragment were used for quantification using an external standard ranging from 100 pM to 10 µM.

### Statistical analysis for bacterial response to nutrient supplementation in filtered seawater

For a given filtered seawater sample with or without nutrient supplementation, multiple replicate experiments were done with single-cell masses and growth rates in each. Rather than ignoring replicate experiments by pooling single cell data across them, statistical significance was assessed by a linear mixed model that considers independent replicate experiments, as previously reported (*47*, *48*). This approach involved pairwise comparisons between two groups of data, each including all replicate experiments, to discern differences attributable to both random and fixed effects. The actual biological effect between the two groups (e.g. nutrient amendment) was considered as fixed effects, and the replicate-to-replicate variation caused by systematic and stochastic fluctuation was considered as random effects. To determine statistical significance, two models were considered: a model as described above and a null model without the fixed effects. Maximum likelihood was calculated for the two models, *L*_*model*_ and *L*_*null*_. The likelihood ratio denotes the degree to which the fitting of the model improves when incorporating a fixed effect, relative to the null model that does not consider it. By applying Wilks’ theorem, which states that the distribution of −2 log(*L*_*null*_⁄*L*_*model*_) approaches a χ^2^ distribution, corresponding p-values were derived.

### Statistical analysis for the nutrient growth law and cell sorting validation

The nutrient growth law analysis was performed on seawater and culture media datasets. In culture media, there were 11 *Vibrionaceae* strains representing 5 species and grown in 9 different media (n = 38 independent strain/media combinations). For each of these 38 unique data points, multiple biological replicate cultures were performed and in each replicate the population growth rate and median mass of cells in the culture was determined. We then calculated the mean of the replicate population growth rate estimates and the mean of the replicate median mass estimates to yield the 38 independent strain/media combinations. Similarly for the seawater dataset (n = 25 independent day/amendment seawater sample combinations), multiple biological replicate cultivation experiments were performed in the SMR and in each replicate the mean growth rate of all cells and the median buoyant mass of cells was determined. We then calculated the mean of all replicate mean growth rate estimates and the mean of the replicate median mass estimates to yield the 25 independent day/amendment seawater sample combinations. First, the culture media and seawater datasets were analyzed separately by fitting an OLS regression model with log10-transformed mean of median mass as the response variable and mean of mean growth rate as the explanatory variable using the R statistical software package. Second, a combined dataset of both culture media and seawater was analyzed with log10-transformed mean of median mass as the response variable and with sample-type (seawater or culture-media), growth rate, and the interaction between growth rate and sample-type included as explanatory variables in an OLS multiple-regression model using the R statistical software package. The growth rate term and intercept were statistically significant (*p* < 2×10^-16^) while the sample-type and interaction term were not statistically significant (*p* = 0.349 and *p* = 0.582, respectively). Since the sample-type and interaction term estimates were not statistically significant, we fit an OLS regression model to the combined dataset with log10-transformed mean of median mass as the response variable and mean of mean growth rate as the only explanatory variable using the R statistical software package. This combined dataset regression without sample-type or the interaction term is plotted in panel 4B as a black line with gray confidence band representing the 95% confidence interval of the regression. We also performed all of the above analyses with ranged major axis regression, which can account for residual error in both variables whereas OLS regression only considers residual variation in the response variable, but the results did not substantially differ from OLS regression. Since none of our conclusions would change by using this more complicated statistical procedure, we do not report them here and include only the OLS results for simplicity.

To assess the mass loss of FACS sorted cells, the mass distributions of 3 replicate *V. cyclitrophicus* 1G07 starved cultures were measured while living (live treatment) and after being fixed, FISH stained, and FACS sorted (sorted treatment). The median mass of each replicate was determined and then an analysis of variance was performed using median mass as the response variable and the treatment (live or sorted) was the explanatory variable in the R statistical software. The difference in mass between the two treatments (in femtograms and as a percent of mass lost from living cells) and statistical significance of this analysis are reported. We then compared the mass of wild *Vibrionaceae* cells after sorting to the mass of starved *V. cyclitrophicus* 1G07 cells after sorting using an analysis of variance with median mass as the response variable and the treatment (wild or starved) was the explanatory variable in the R statistical software.

## Acknowledgement

We would like to acknowledge many past lab members from our groups, collaborators, and friends for help on many aspects of this work. Kathryn Kauffman for the 2010 Nahant seawater samples and providing advice for our 2023 sampling. Fatima Hussain, Joseph Elsherbini, Kathryn Kauffman, and Xiaoqian (Annie) Yu for isolating strains from Nahant seawater used in this study. Dan Distel, Rosie Falco, Ella Messner, Yejin Hwang, Rachel Szabo, Gabriel Vercelli, Rachel Gregor, Otto Cordero, Stefany Moreno, Samantha Pennino, M.S. Suryateja Jammalamadaka, Chris Sedlacek, and Kathi Kitzinger for assistance organizing, sampling, and processing seawater in 2023. We would also like to acknowledge the Life Science Compute Cluster (LISC) of the University of Vienna which was used to achieve the raw processing of the amplicon sequencing data presented in this work. This work was funded by the following: The Simons Foundation (Life Sciences Project Award 572792 to M.F.P. and S.R.M.); the National Science Foundation (Award 2319028 to S.R.M.); and Austrian Science Fund (FWF Grant DOI: 10.55776/COE7 to M.F.P.).

---

**Fig. S1.**
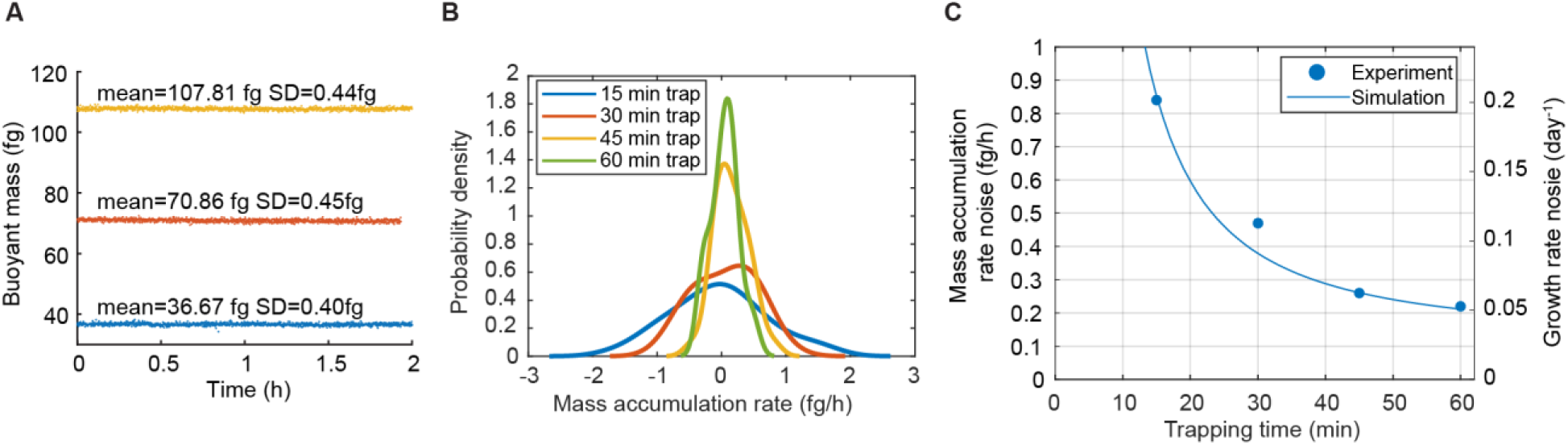
Resolution of SMR mass and growth rate measurement. **(A)** Continuous buoyant mass measurements by trapping of polystyrene microbeads with the diameter of 1.1, 1.3, and 1.6 µm. **(B)** Mass accumulation rate distribution of 1.3 µm microbeads trapped for different time periods. **(C)** Noise level of mass accumulation rate and growth rate in relation to different trapping times. The noise of mass accumulation rate was experimentally obtained from (B) (blue dots) and simulated as the standard error of slope derived from linear regression—a function of mass error in (A) and trapping time (blue lines). The mass accumulation rate on the left axis was converted to the growth rate on the right axis assuming a buoyant mass of 100 fg.

**Fig. S2.**
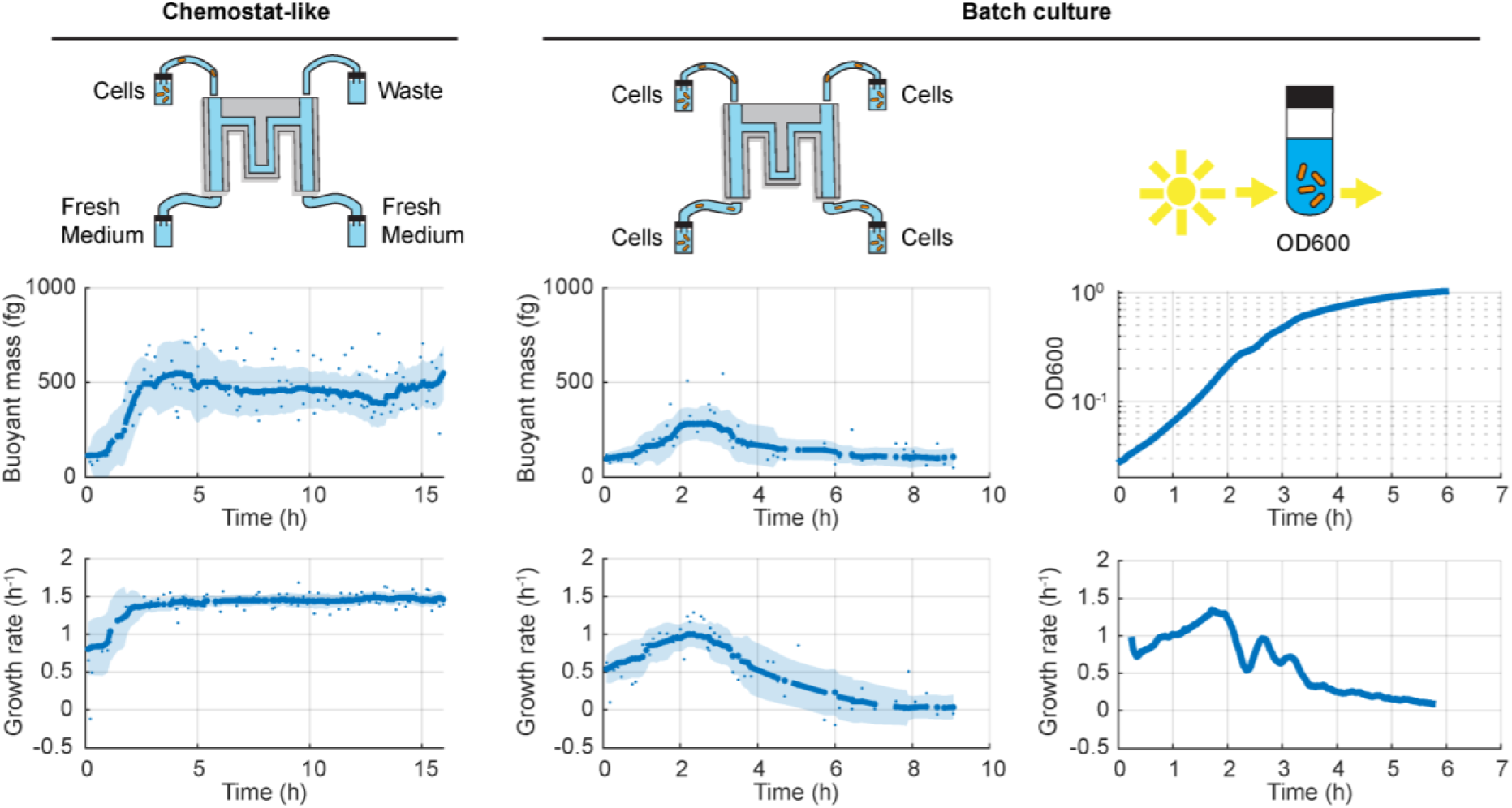
SMR mimics a miniaturized chemostat and maintains steady-state mass and growth rate profiles in typical laboratory media. The physiologically stable, chemostat-like environment o SMR trapping system was comparatively illustrated by *Vibrio cyclitrophicus* growing in Marine Broth 2216 under three experimental settings. (**Left**) Cell-free fresh medium is continuously provided to the bacteria, removing any effects of nutrient depletion on growth. (**Middle**) All vials of the SMR device contain cells without fresh medium input over time, mimicking a batch culture. (**Right**) A separate batch culture in a cell culture tube with real-time OD600 measurements. The growth in the two batch cultures (middle and right) slowed down after 2 h and dropped to almost zero after 6 h, due to onset of nutrient depletion in batch culture. On the contrary, the growth rate and cell mass in the chemostat (left) were maintained at the maximum for a prolonged period. The dots in the left and middle panels refer to single-cell measurements. The lines in the left and middle panels refer to the moving averages of the corresponding dots, and the shades refer to the standard deviation of the moving averaging.

**Fig. S3.**
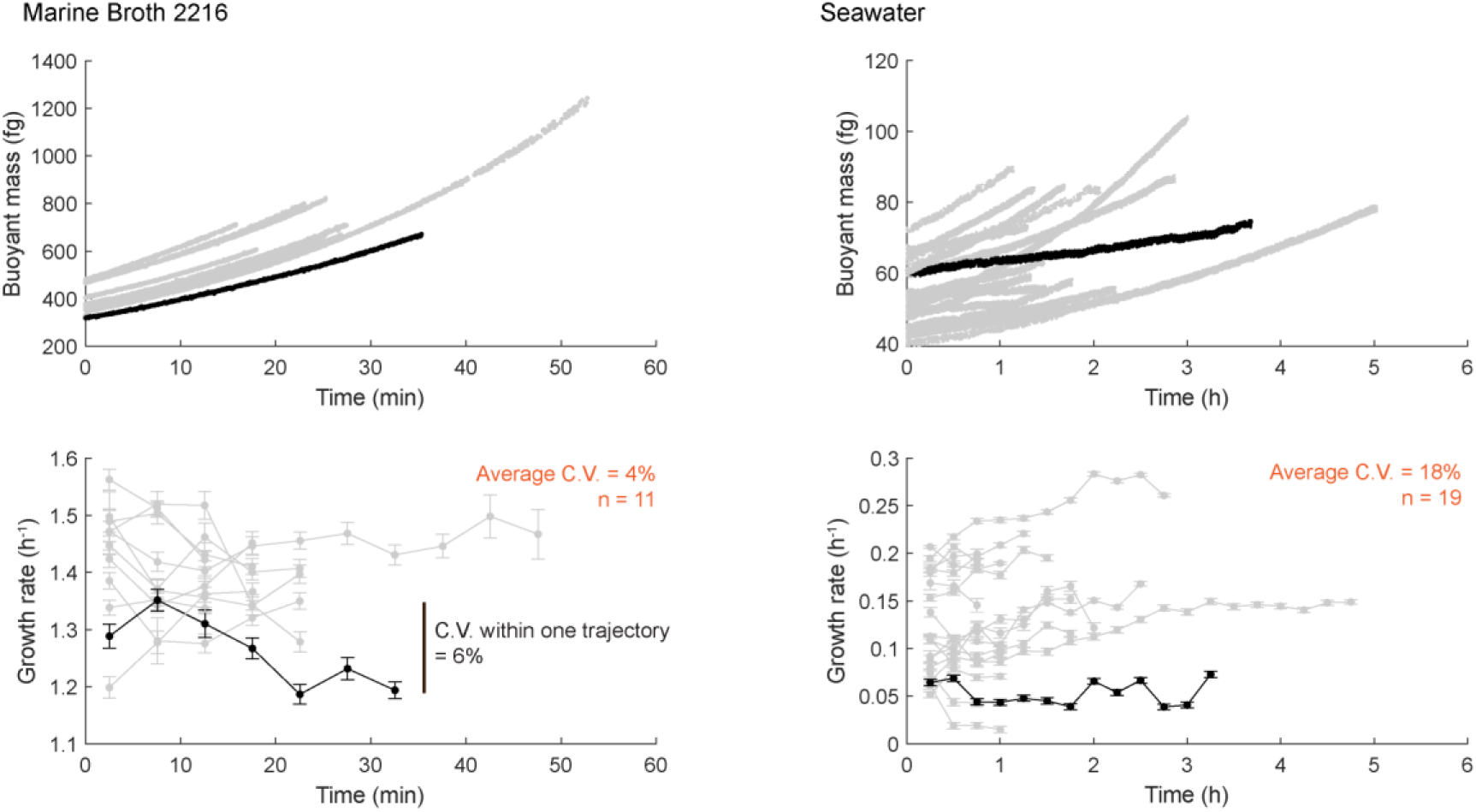
Single-cell growth trajectories of *Vibrio cyclitrophicus* in Marine Broth 2216 and filtered seawater. Each trajectory was longer than the set trapping time of 5 min (fast mode) or 30 min (slow mode) to evaluate trapping time-dependent variation in results. For each trajectory, instantaneous growth rates were calculated from bins with the width of 5 or 30 min, and the coefficient of variation (C.V.) of growth rate was calculated. The growth rate C.V. of all trajectories from the two conditions was averaged, respectively.

**Fig. S4.**
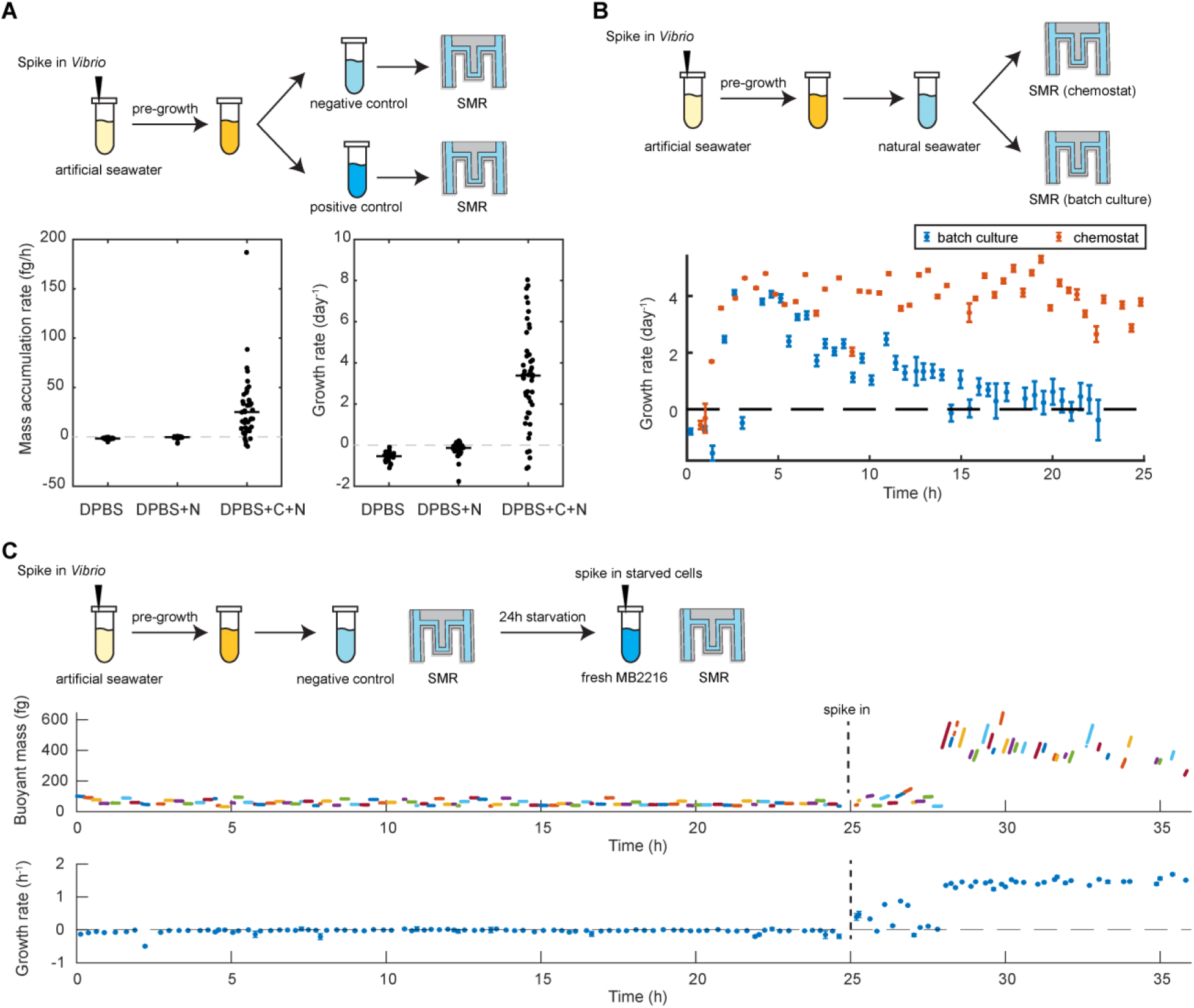
Negative and positive controls of *Vibrio* growth in the SMR system. **(A)** Parallel comparison between negative and positive controls. *Vibrio cyclitrophicus* bacteria were pre-grown into stationary phase and separately transferred to Dulbecco’s Phosphate-Buffered Saline (DPBS), nitrogen-amended DPBS (DPBS+N), and nitrogen and carbon-amended DPBS (DPBS+C+N). Carbon was added as 10 mM glucose and nitrogen was added as 10 mM NH_4_Cl. **(B)** Nutrients in filtered seawater could be fully consumed in the batch culture mode. *Vibrio cyclitrophicus* bacteria were pre-grown into stationary phase and transferred to unamended filtered seawater. In addition to running the SMR in the chemostat mode where cells were loaded into one vial, the measurement was repeated in the batch culture mode with cells in all vials to reveal the full consumption of nutrients in the seawater. **(C)** Sequential comparison between negative and positive controls. *Vibrio cyclitrophicus* bacteria were pre-grown into stationary phase and transferred to Phosphate-Buffered Saline (PBS) as a negative control, followed by transfer to Marine Broth 2216 as a positive control.

**Fig. S5.**
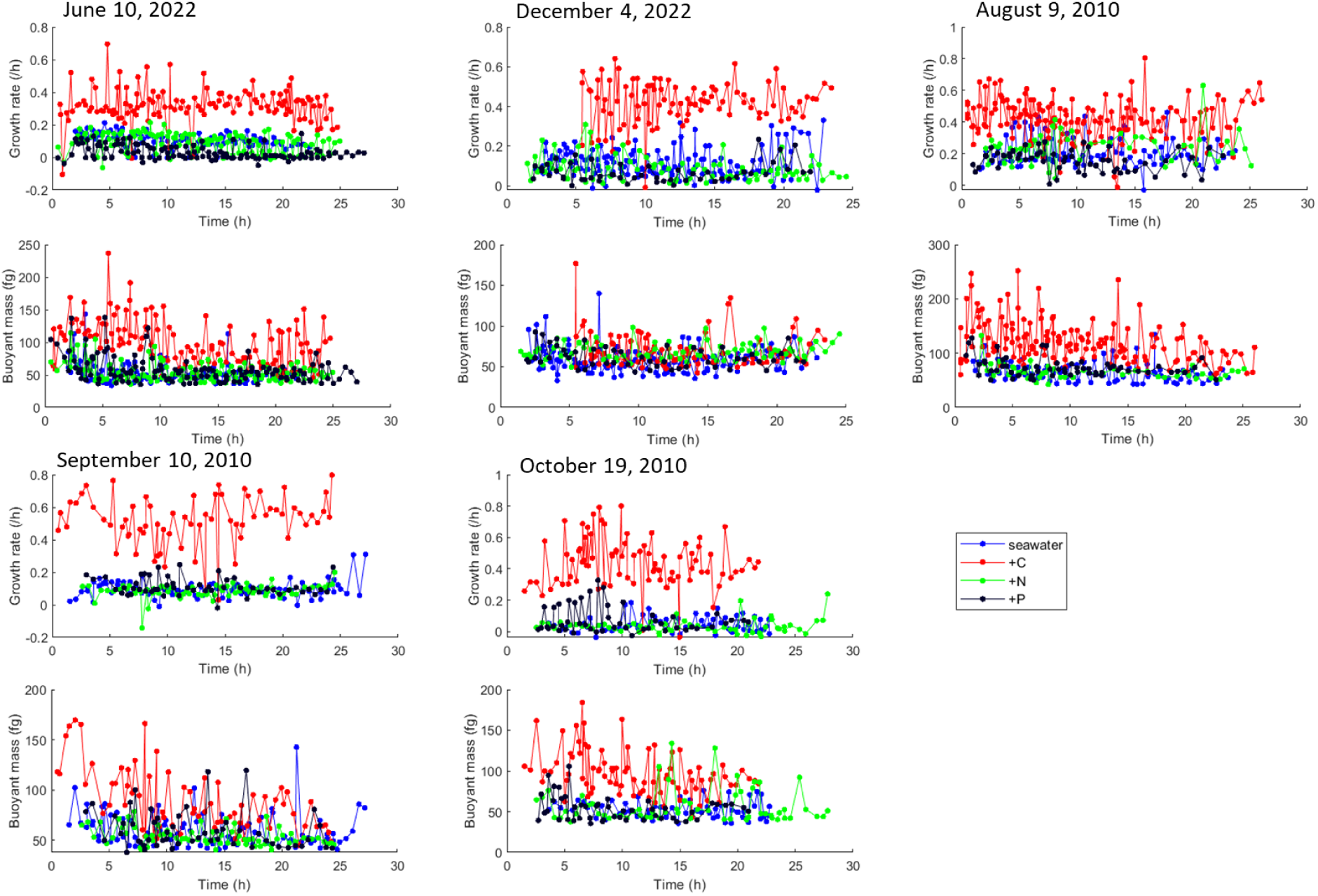
Time-lapse growth rate and buoyant mass of *Vibrio cyclitrophicus* reaching steady states in seawater. Each dot refers to a single-cell measurement and those from the same condition (with replicates combined) are connected in a line. The acclimation period (first 1–3 hours) is removed from each condition.

**Fig. S6.**
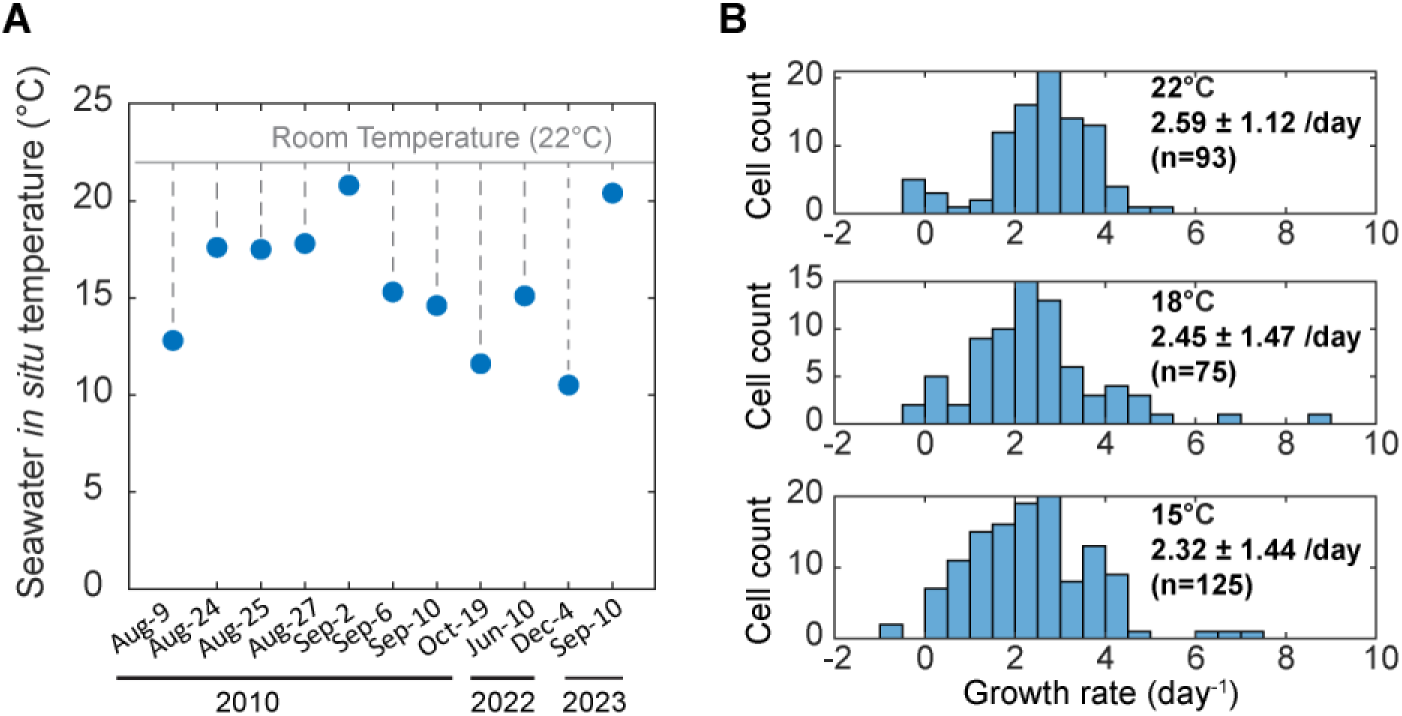
Growth rate of *Vibrio cyclitrophicus* is not significantly altered when measured at room temperature versus *in situ* temperatures. **(A)** Seawater *in situ* temperatures vary across seawater samples, some of which are close to the room temperature (22°C) at which the SMR experiments were done. **(B)** Growth rate distribution of *V. cyclitrophicus* in the seawater sample collected on June 10, 2022, measured at three different temperatures.

**Fig. S7.**
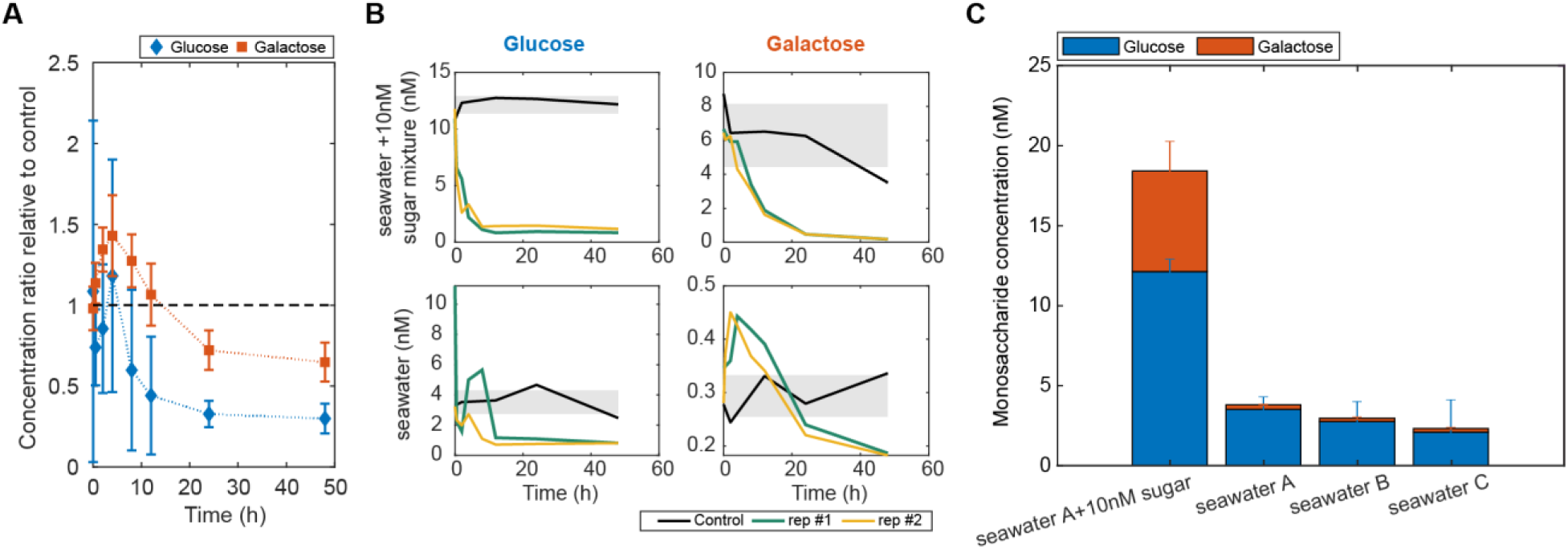
Metabolite uptake of *Vibrio cyclitrophicus* in filtered seawater. **(A)** Glucose and galactose concentrations normalized by the initial, unspent concentration (calculated as the average concentration of the corresponding negative control) and averaged across three filtered seawater samples. **(B)** Glucose and galactose concentrations over time in the supernatant of *V. cyclitrophicus* cultures in a seawater sample collected on June 10, 2022, either unamended or amended with a mixture of monosaccharides (fucose, galactose, glucosamine, glucose, mannose, rhamnose, xylose; 10nM each). **(C)** The initial, unspent concentrations of glucose and galactose in four seawater conditions.

**Fig. S8.**
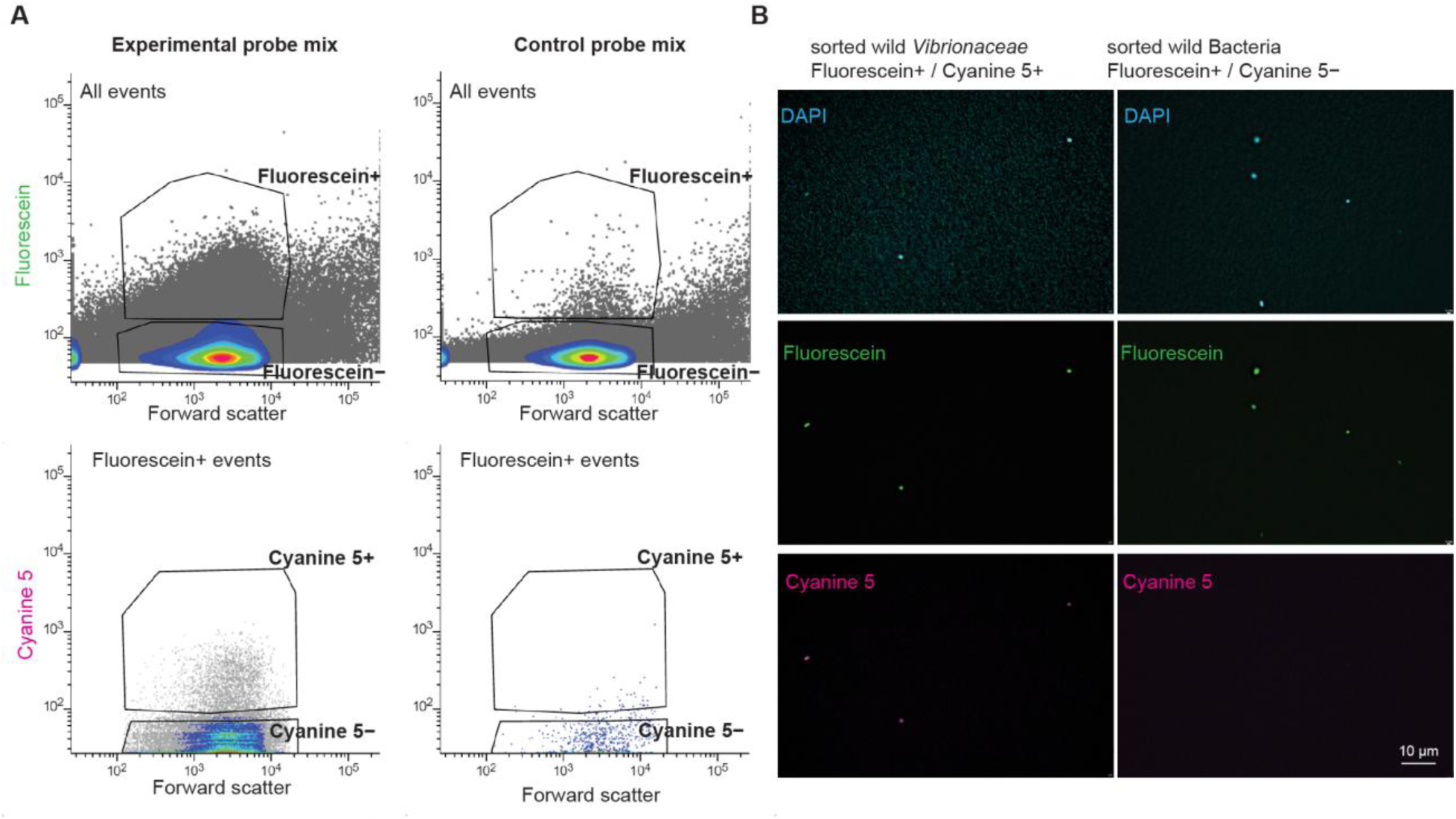
FISH labeling and FACS sorting of environmental samples. **(A)** Cells were dual-labeled with two probe mixes (experimental or control) each containing two FISH probes attached to different fluorophores (Fluorescein or Cy5), then sorted based on these two fluorescence phenotypes. In experimental probe mix, double positive sorted cells are Sorted Vibrio, while Fluorescein positive and Cy5 negative sorted cells are Sorted Bacteria. In control probe mix, double positive sorted cells are False Positive, while double negative sorted cells are True Negative. **(B)** Imaging of sorted cells demonstrates FISH labeling and FACS sorting of seawater samples. Sorted Vibrio cells and Sorted Bacteria cells were imaged with fluorescence microscopy on 0.2µm PC membrane, with representative images from a field of view from replicate 3 shown. DAPI was used to stain nucleic acids in addition to the FISH probes. All DAPI stained cells showed expected fluorescence phenotypes for each sorted population (3 cells in Sorted Vibrio, 5 cells in Sorted Bacteria).

**Fig. S9.**
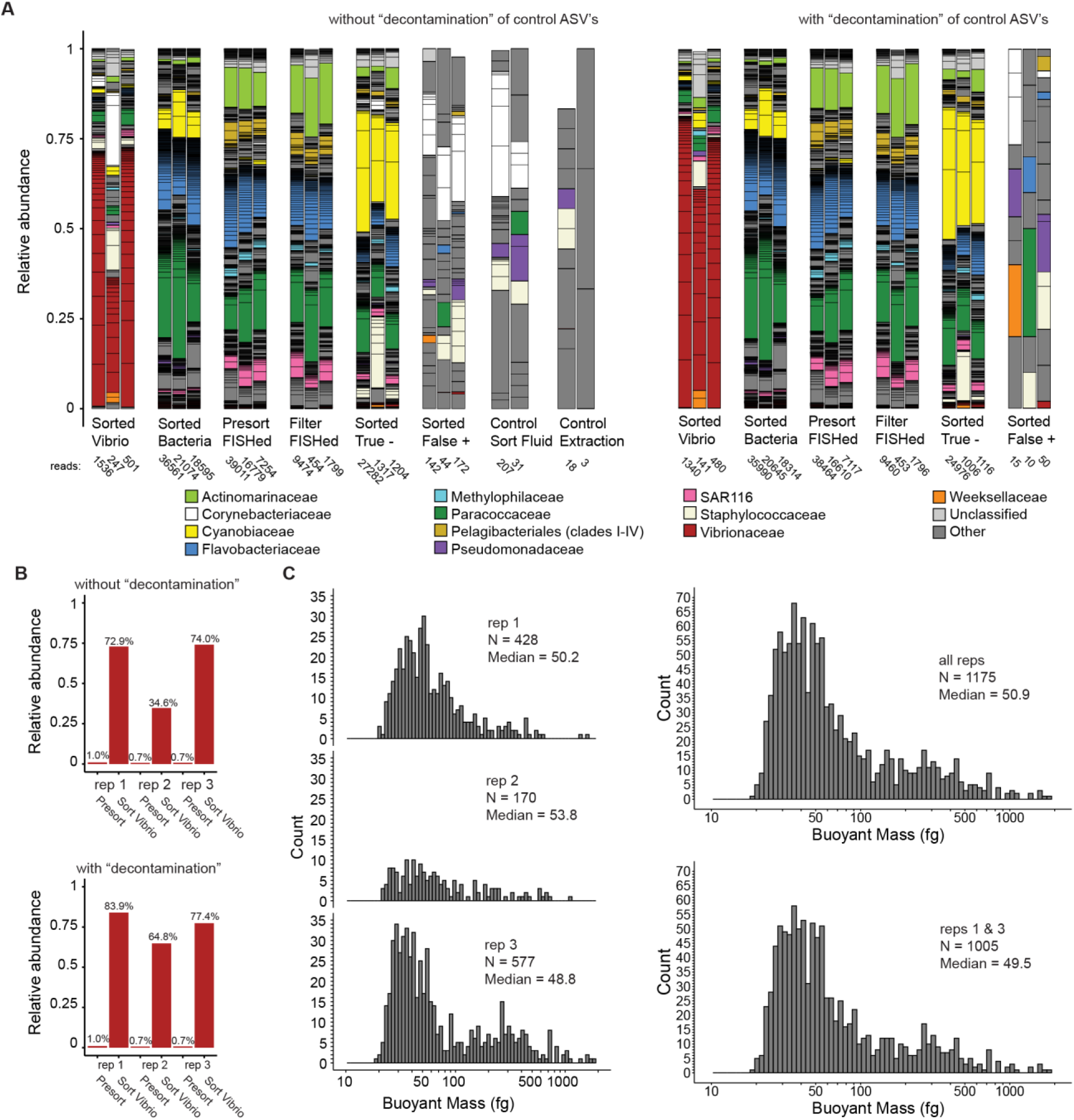
Validation of FISH-labeled cell sorting and subsequent mass measurements. **(A)** 16S rRNA amplicon sequencing validates successful FISH labeling and FACS sorting of seawater samples. All sorted cell populations, filter samples, and control samples were sequenced and are presented with decontamination (removing ASV sequences found in control samples from all other samples) or without decontamination. All seawater samples have 3 replicates (plotted left to right 1-3) collected from different spatial locations within our sampling site, while controls of sorting fluid has two replicates from separate days of sorting and two DNA extraction kits were used to extract different samples. Decontamination had minimal impact on seawater samples, but led to a higher relative abundance of *Vibrionaceae* sequences in sorted Vibrio populations. **(B)** *Vibrionaceae* sequences were enriched with or without decontamination. Notably, the *Vibrionaceae* in decontaminated sequences showed an increase in average relative abundance from 0.8% of the Presort community to 75.3% of the Sorted Vibrio sample. **(C)** Mass distributions for individual replicates of Sorted Vibrio populations show consistent medians across all replicates, but fewer cells collected from replicate 2. The median mass of all combined replicates, or for only replicates 1 & 3 do not differ substantially.

**Fig. S10.**
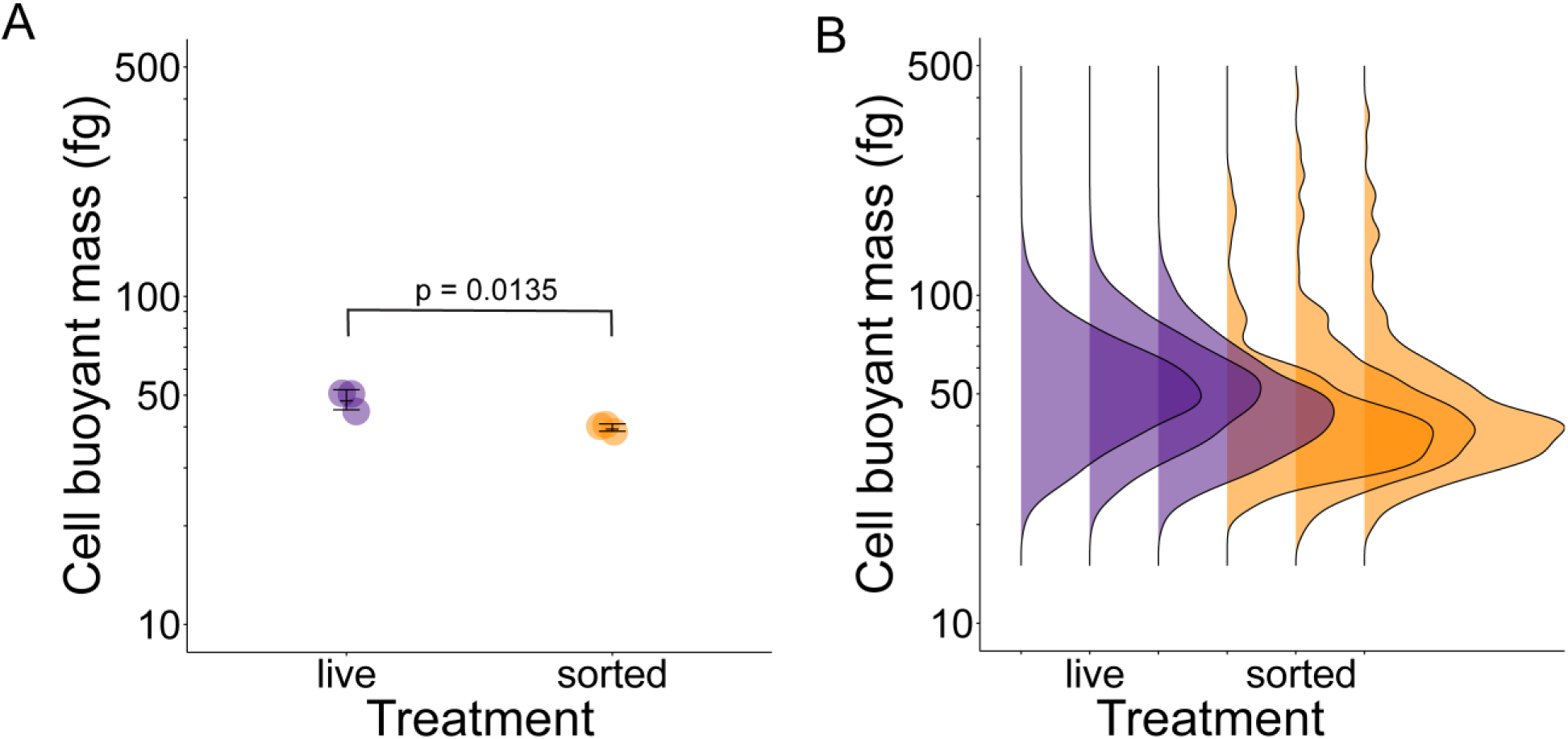
Mass alteration of starved *Vibrio cyclitrophicus* cells due to fixation procedures. The average cell mass **(A)** (n = 3 replicate cultures each) and cell mass distributions **(B)** (N= 6925, 6802, 9296 cells in live populations; N = 450, 393, 366 cells in sorted populations) of *V. cyclitrophicus* cultures starved for 20 hours in PBS. Live cells have significantly larger mass than sorted cells (8.72 fg mass difference or 17.96% mass loss, *p* = 0.0135) after experiencing all sample processing steps (filtration, freezing, FISH, & FACS).

**Fig. S11.**
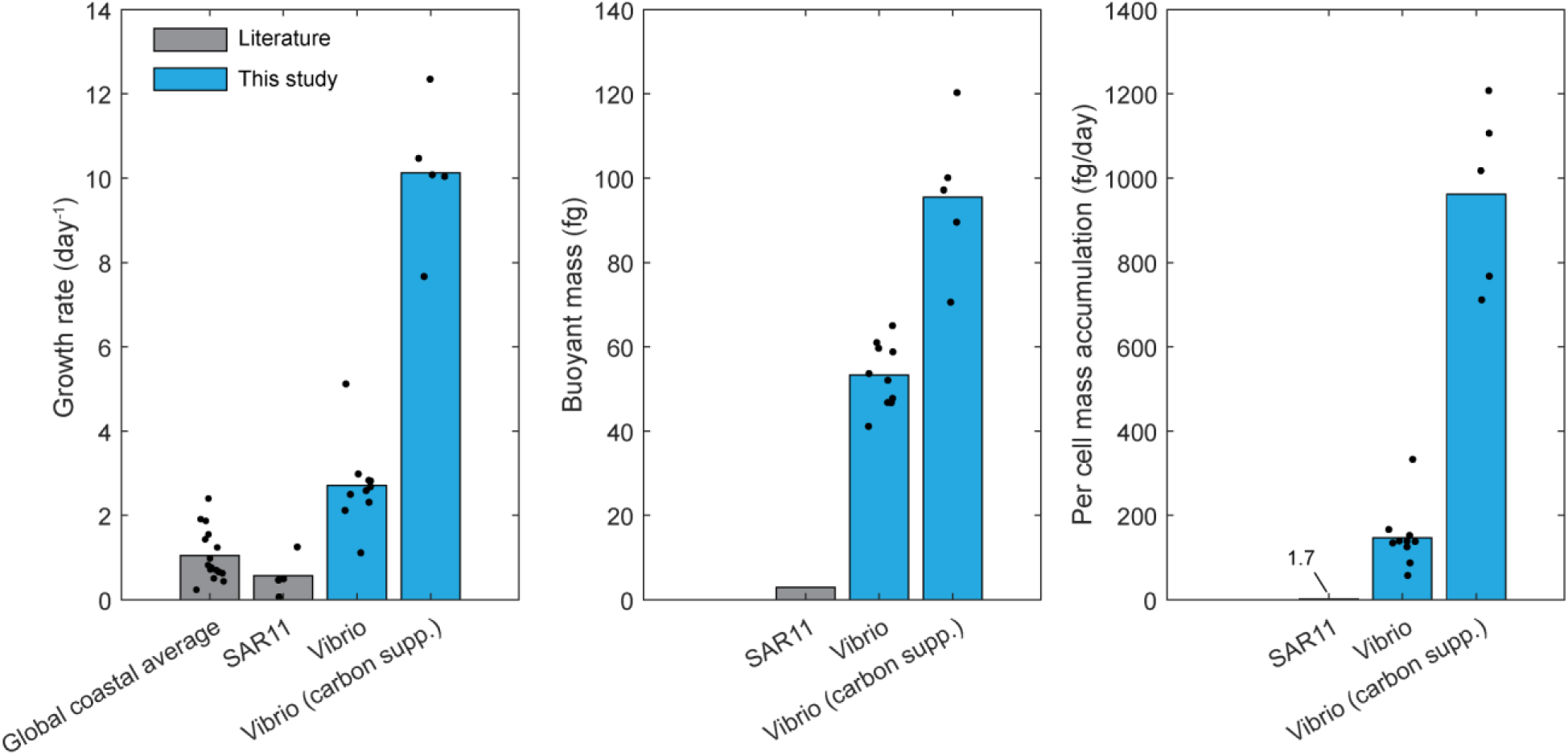
Growth comparison of *Vibrio* in this study and literature results of whole bacterial community and SAR11 clade in global coastal environments. For both literature and results from this study, each dot refers to a measurement of whole community or certain taxa from either global coastal samples or our Nahant time-series samples, and each bar refers to the averages. The complete references for the literature results are listed in table S6.

## References

1. J. S. Poindexter, “Oligotrophy: fast and famine existence” in Advances in Microbial Ecology, M. Alexander, Ed. (Springer US, Boston, MA, 1981; http://link.springer.com/10.1007/978-1-4615-8306-6_2)vol. 5, pp. 63–89.

2. D. L. Kirchman, Growth Rates of Microbes in the Oceans. Annu. Rev. Mar. Sci. 8, 285–309 (2016).

3. A. Ho, D. P. D. Lonardo, P. L. E. Bodelier, Revisiting life strategy concepts in environmental microbial ecology. Microbiology Ecology, fix006 (2017).

4. J. R. Thompson, M. F. Polz, “Dynamics of *Vibrio* Populations and Their Role in Environmental Nutrient Cycling” in The Biology of Vibrios, F. L. Thompson, B. Austin, J. Swings, Eds. (ASM Press, Washington, DC, USA, 2006; http://doi.wiley.com/10.1128/9781555815714.ch13), pp. 190–203.

5. J. H. Munson-McGee, M. R. Lindsay, E. Sintes, J. M. Brown, T. D’Angelo, J. Brown, L. C. Lubelczyk, P. Tomko, D. Emerson, B. N. Orcutt, N. J. Poulton, G. J. Herndl, R. Stepanauskas, Decoupling of respiration rates and abundance in marine prokaryoplankton. Nature 612, 764–770 (2022).

6. J. D. Brüwer, L. H. Orellana, C. Sidhu, H. C. L. Klip, C. L. Meunier, M. Boersma, K. H. Wiltshire, R. Amann, B. M. Fuchs, In situ cell division and mortality rates of SAR11, SAR86, Bacteroidetes, and Aurantivirga during phytoplankton blooms reveal differences in population controls. mSystems, e0128722 (2023).

7. O. Deulofeu-Capo, M. Sebastián, A. Auladell, C. Cardelús, I. Ferrera, O. Sánchez, J. M. Gasol, Growth rates of marine prokaryotes are extremely diverse, even among closely related taxa. ISME Communications 4, ycae066 (2024).

8. L. K. Fecskeová, K. Piwosz, D. Šantić, S. Šestanović, A. V. Tomaš, M. Hanusová, M. Šolić, M. Koblížek, Lineage-Specific Growth Curves Document Large Differences in Response of Individual Groups of Marine Bacteria to the Top-Down and Bottom-Up Controls. mSystems 6, 10.1128/msystems.00934-21 (2021).

9. T. P. Burg, M. Godin, S. M. Knudsen, W. Shen, G. Carlson, J. S. Foster, K. Babcock, S. R. Manalis, Weighing of biomolecules, single cells and single nanoparticles in fluid. Nature 446, 1066–1069 (2007).

10. M. Godin, F. F. Delgado, S. Son, W. H. Grover, A. K. Bryan, A. Tzur, P. Jorgensen, K. Payer, A. D. Grossman, M. W. Kirschner, S. R. Manalis, Using buoyant mass to measure the growth of single cells. Nat Methods 7, 387–390 (2010).

11. E. Levien, J. H. Kang, K. Biswas, S. R. Manalis, A. Amir, T. P. Miettinen, Stochasticity in mammalian cell growth rates drives cell-to-cell variability independently of cell size and divisions. Cold Spring Harbor Laboratory [Preprint] (2025). 10.1101/2025.06.18.659700.

12. B. J. Shapiro, J. Friedman, O. X. Cordero, S. P. Preheim, S. C. Timberlake, G. Szabó, M. F. Polz, E. J. Alm, Population Genomics of Early Events in the Ecological Differentiation of Bacteria. Science 336, 48–51 (2012).

13. Y. Yawata, O. X. Cordero, F. Menolascina, J.-H. Hehemann, M. F. Polz, R. Stocker, Competition– dispersal tradeoff ecologically differentiates recently speciated marine bacterioplankton populations. Proc. Natl. Acad. Sci. U.S.A. 111, 5622–5627 (2014).

14. N. Cermak, J. W. Becker, S. M. Knudsen, S. W. Chisholm, S. R. Manalis, M. F. Polz, Direct single-cell biomass estimates for marine bacteria via Archimedes’ principle. ISME J 11, 825–828 (2016).

15. T. P. Miettinen, K. S. Ly, A. Lam, S. R. Manalis, Single-cell monitoring of dry mass and dry mass density reveals exocytosis of cellular dry contents in mitosis. eLife 11, e76664 (2022).

16. B. R. K. Roller, C. Hellerschmied, Y. Wu, T. P. Miettinen, A. L. Gomez, S. R. Manalis, M. F. Polz, Single-cell mass distributions reveal simple rules for achieving steady-state growth. mBio 14, e01585–23 (2023).

17. S. Giovannoni, U. Stingl, The importance of culturing bacterioplankton in the “omics” age. Nat Rev Microbiol 5, 820–826 (2007).

18. M. J. Church, “Resource Control of Bacterial Dynamics in the Sea” in Microbial Ecology of the Oceans, D. L. Kirchman, Ed. (Wiley, ed. 1, 2008; https://onlinelibrary.wiley.com/doi/10.1002/9780470281840.ch10), pp. 335–382.

19. M. Sebastián, J. M. Gasol, Heterogeneity in the nutrient limitation of different bacterioplankton groups in the Eastern Mediterranean Sea. ISME J 7, 1665–1668 (2013).

20. Hawaii Ocean Time-series (HOT) Dissolved Organic Matter. https://hahana.soest.hawaii.edu/hot/methods/onuts.html.

21. Hawaii Ocean Time-series (HOT) Dissolved Inorganic Nutrients. https://hahana.soest.hawaii.edu/hot/methods/inuts.html.

22. Winter dissolved inorganic nitrogen (NH4 + NO3 + NO2) concentration observed in European seas, 2013-2017. https://www.eea.europa.eu/data-and-maps/figures/winter-dissolved-inorganic-nitrogen-ammonium-1.

23. Annual mean total phosphorus concentrations in European seas, 2013-2017. https://www.eea.europa.eu/data-and-maps/figures/annual-mean-total-phosphorus-concentrations.

24. A. M. Martin-Platero, B. Cleary, K. Kauffman, S. P. Preheim, D. J. McGillicuddy, E. J. Alm, M. F. Polz, High resolution time series reveals cohesive but short-lived communities in coastal plankton. Nat Commun 9, 266 (2018).

25. M. Schaechter, O. MaalOe, N. O. Kjeldgaard, Dependency on Medium and Temperature of Cell Size and Chemical Composition during Balanced Growth of Salmonella typhimurium. Journal of General Microbiology 19, 592–606 (1958).

26. S. Taheri-Araghi, S. D. Brown, J. T. Sauls, D. B. McIntosh, S. Jun, Single-Cell Physiology. Annu. Rev. Biophys. 44, 123–142 (2015).

27. S. Taheri-Araghi, S. Bradde, J. T. Sauls, N. S. Hill, P. A. Levin, J. Paulsson, M. Vergassola, S. Jun, Cell-Size Control and Homeostasis in Bacteria. Current Biology 25, 385–391 (2015).

28. H. Zheng, Y. Bai, M. Jiang, T. A. Tokuyasu, X. Huang, F. Zhong, Y. Wu, X. Fu, N. Kleckner, T. Hwa, C. Liu, General quantitative relations linking cell growth and the cell cycle in Escherichia coli. Nat Microbiol 5, 995–1001 (2020).

29. S. Vadia, P. A. Levin, Growth rate and cell size: a re-examination of the growth law. Current Opinion in Microbiology 24, 96–103 (2015).

30. P. A. Del Giorgio, J. M. Gasol, D. Vaqué, P. Mura, S. Agustí, C. M. Duarte, Bacterioplankton community structure: Protists control net production and the proportion of active bacteria in a coastal marine community. Limnology & Oceanography 41, 1169–1179 (1996).

31. M. Scott, C. W. Gunderson, E. M. Mateescu, Z. Zhang, T. Hwa, Interdependence of Cell Growth and Gene Expression: Origins and Consequences. Science 330, 1099–1102 (2010).

32. J. Lee, R. Chunara, W. Shen, K. Payer, K. Babcock, T. P. Burg, S. R. Manalis, Suspended microchannel resonators with piezoresistive sensors. Lab Chip 11, 645–651 (2011).

33. K. M. Kauffman, W. K. Chang, J. M. Brown, F. A. Hussain, J. Yang, M. F. Polz, L. Kelly, Resolving the structure of phage–bacteria interactions in the context of natural diversity. Nat Commun 13, 372 (2022).

34. X. A. Yu, C. McLean, J.-H. Hehemann, D. Angeles-Albores, F. Wu, A. Muszyński, C. H. Corzett, P. Azadi, E. B. Kujawinski, E. J. Alm, M. F. Polz, Low-level resource partitioning supports coexistence among functionally redundant bacteria during successional dynamics. The ISME Journal 18, wrad013 (2024).

35. H. Daims, Use of Fluorescence In Situ Hybridization and the *daime* Image Analysis Program for the Cultivation-Independent Quantification of Microorganisms in Environmental and Medical Samples. Cold Spring Harb Protoc 2009, pdb.prot5253 (2009).

36. R. Sekar, B. M. Fuchs, R. Amann, J. Pernthaler, Flow Sorting of Marine Bacterioplankton after Fluorescence In Situ Hybridization. Appl Environ Microbiol 70, 6210–6219 (2004).

37. G. Wallner, R. Amann, W. Beisker, Optimizing fluorescent in situ hybridization with rRNA-targeted oligonucleotide probes for flow cytometric identification of microorganisms. Cytometry 14, 136–143 (1993).

38. R. I. Amann, B. J. Binder, R. J. Olson, S. W. Chisholm, R. Devereux, D. A. Stahl, Combination of 16S rRNA-targeted oligonucleotide probes with flow cytometry for analyzing mixed microbial populations. Appl Environ Microbiol 56, 1919–1925 (1990).

39. L. Girard, S. Peuchet, P. Servais, A. Henry, N. Charni-Ben-Tabassi, J. Baudart, Spatiotemporal Dynamics of Total Viable *Vibrio* spp. in a NW Mediterranean Coastal Area. Microbes and environments 32, 210–218 (2017).

40. A. Klindworth, E. Pruesse, T. Schweer, J. Peplies, C. Quast, M. Horn, F. O. Glöckner, Evaluation of general 16S ribosomal RNA gene PCR primers for classical and next-generation sequencing-based diversity studies. Nucleic Acids Research 41, e1–e1 (2013).

41. P. Pjevac, B. Hausmann, J. Schwarz, G. Kohl, C. W. Herbold, A. Loy, D. Berry, An Economical and Flexible Dual Barcoding, Two-Step PCR Approach for Highly Multiplexed Amplicon Sequencing. Front. Microbiol. 12, 669776 (2021).

42. B. Langmead, S. L. Salzberg, Fast gapped-read alignment with Bowtie 2. Nat Methods 9, 357–359 (2012).

43. D. E. Wood, J. Lu, B. Langmead, Improved metagenomic analysis with Kraken 2. Genome Biol 20, 257 (2019).

44. P. Danecek, J. K. Bonfield, J. Liddle, J. Marshall, V. Ohan, M. O. Pollard, A. Whitwham, T. Keane, S. A. McCarthy, R. M. Davies, H. Li, Twelve years of SAMtools and BCFtools. GigaScience 10, giab008 (2021).

45. B. Rühmann, J. Schmid, V. Sieber, Fast carbohydrate analysis via liquid chromatography coupled with ultra violet and electrospray ionization ion trap detection in 96-well format. Journal of Chromatography A 1350, 44–50 (2014).

46. G. Xu, M. J. Amicucci, Z. Cheng, A. G. Galermo, C. B. Lebrilla, Revisiting monosaccharide analysis – quantitation of a comprehensive set of monosaccharides using dynamic multiple reaction monitoring. Analyst 143, 200–207 (2018).

47. M. Herbig, M. Kräter, K. Plak, P. Müller, J. Guck, O. Otto, “Real-Time Deformability Cytometry: Label-Free Functional Characterization of Cells” in Flow Cytometry Protocols, T. S. Hawley, R. G. Hawley, Eds. (Springer New York, New York, NY, 2018; http://link.springer.com/10.1007/978-1-4939-7346-0_15)vol. 1678 of *Methods in Molecular Biology*, pp. 347–369.

48. M. Herbig, A. Mietke, P. Müller, O. Otto, Statistics for real-time deformability cytometry: Clustering, dimensionality reduction, and significance testing. Biomicrofluidics 12, 042214 (2018).

