## supplementary text for "Single-cell mass accumulation reveals bacterioplankton growth rate in native seawater"

### Supplementary Note #1: Technical characterization and validation of SMR growth measurement

**Measurement resolution**

We first evaluated the resolution of mass measurement by repeatedly measuring a polystyrene microbead for hours. Three polystyrene microbeads (1.1, 1.3, and 1.6 μm in diameter) were measured every ~10 seconds for 2 hours, respectively, providing repeated measurements of the same microbead whose standard deviation indicates mass resolution. The three microbeads covered a wide range of buoyant mass from 36.7 to 107.8 fg and showed narrow variations within the 2 hours indicating mass resolutions of 0.4 – 0.45 fg. (**fig. S1A**). Then we characterized the resolution of growth rate measurement by measuring a 1.3-μm microbead population following the same cell trapping procedures described for bacteria in the Method section. The polystyrene beads were considered to have zero growth rate as ground truth and the magnitude of deviation from the true zero constitutes a measure of growth rate resolution. The resolution was characterized at different trapping times including 15, 30, 45, and 60 minutes. We found that as trapping time decreased from 15 to 60 minutes, growth rates were more narrowly distributed around the true zero and the resolutions of mass accumulation rate and specific growth rate became finer (**fig. S1B**).

Since mass accumulation rate is intrinsically defined by the linear regression of mass vs time, its resolution can also be estimated by the standard error of the linear regression slope coefficient estimator when assuming the error term of mass. For a linear regression model of $y = \alpha + \beta x + \varepsilon$ between variables $y$ and $x$, the estimator of slope coefficient $\beta$ will be normally distributed with variance $\sigma^2/\sum(x_i - \overline{x})^2$ where $\sigma$ is the standard deviation of the error term. Hence, the standard error of $\beta$, $s_\beta$,can be described as follows:

$$s_\beta = \sqrt{\frac{\sigma^2}{\sum(x_i - \overline{x})^2}}$$

In our setup, $x_i$ is a series of uniformly spaced time points, for example $x_i$ = 0s, 5s, 10s, 15s… Define the total number of measurements as $n$, the time increment as $\Delta t$, and the total time as $T$ ($T = n \cdot \Delta t$). Then the term $\sum(x_i - \overline{x})^2$ can be re-written as:

$$\sum(x_i - \overline{x})^2 = \Delta t^2 \sum\left(i - \frac{n+1}{2}\right)^2$$

$$= \Delta t^2 \left[\left(\frac{-n+1}{2}\right)^2 + \left(\frac{-n+3}{2}\right)^2 + \cdots + \left(\frac{n-3}{2}\right)^2 + \left(\frac{n-1}{2}\right)^2\right] = \Delta t^2 \cdot \frac{n^3 - n}{12}$$

Finally, $s_\beta$ can be re-written as follows:

$$s_\beta = \frac{\sqrt{12} \cdot \sigma}{\Delta t \sqrt{n^3 - n}} = \frac{\sqrt{12} \cdot \sigma}{T\sqrt{n - \frac{1}{n}}}$$

When measuring the growth rates of microbead populations, the resolution of mass accumulation rate was simulated as $s_\beta$ assuming a constant standard deviation of mass $\sigma$ = 0.4 fg at different trapping times. The simulated resolution agreed with the experimental results at trapping times of 15 – 60 minutes (**fig. S1C**). Combing both experimental and simulated results, the SMR can resolve buoyant mass by 0.4 fg, and when trapping for 30 minutes, mass accumulation rate by ~0.4 fg/h and grow rate by ~0.1 /day (normalized by a buoyant mass of 100 fg).

**Cell trapping time validation**

In this study, individual bacterial cells were set to be trapped for 5 minutes in laboratory culture media and for 30 minutes in filtered seawater. The measured growth rates usually led to a doubling time longer than the set trapping time. To understand the uncertainty of estimating growth rate from a short duration, we trapped a few cells longer than the usual trapping time in marine broth 2216 and filtered seawater, respectively. The longest trap reached 55 minutes in marine broth 2216 and 5 hours in seawater. Each long trap was broken into short traps of 5 or 30 minutes, and the variation across growth rate estimates from these short traps was calculated. These short-trap growth rates showed a C.V. of 4% on average in marine broth 2216 and 18% in seawater, suggesting that estimating growth from a shorter duration did not substantially distort the measurement (**fig. S3**).

**Biological controls**

To validate that the SMR system does not introduce contaminant to seawater that drives the measured growth in our results, we conducted biological control experiments in the following sense: Firstly, we measured growth in paired negative and positive controls which only differed by the addition of organic carbon and/or inorganic nitrogen. Secondly, we switched to the batch culture mode of the SMR system and showed that the growth-supporting nutrients in seawater were not unlimited and could be depleted over time. Thirdly, we sequentially grew cells in negative control and positive control to demonstrate that the cells in the negative control were viable and their growth could be recovered upon transfer to positive control.

In the first experiment, *Vibrio cyclitrophicus* bacteria were pre-grown in artificial seawater into stationary phase and separately transferred to Dulbecco's Phosphate-Buffered Saline (DPBS), nitrogen-amended DPBS (DBPS+N) and nitrogen and carbon-amended DPBS (DBPS+C+N). Carbon was added as 10 mM glucose and nitrogen was added as 10 mM $NH_4Cl$. The bacteria only grew under DPBS+C+N and did not under either DPBS+N or DPBS, suggesting that there was at least no organic carbon leaching from the SMR system (**fig. S4A**). In the second experiment, *Vibrio cyclitrophicus* bacteria were pre-grown into stationary phase, transferred to unamended seawater, and measured in both chemostat mode (where cells were loaded into one vial) and in batch culture mode (where cells were loaded in all vials). As opposed to continuous growth in the chemostat mode, the growth rate in the batch culture mode decreased after 5 hours and reached zero at ~20 hours, suggesting that the nutrients were depleted over time. If any nutrient leaching existed in the SMR system and caused the observed growth across every independent experiment, one would expect that the "leached nutrients" persist from experiment to experiment and never run out throughout an experiment. This is rejected by the nutrient depletion that we observed in the batch culture mode, proving that the observed growth is not an artifact (**fig. S4B**). In the third experiment, *Vibrio cyclitrophicus* bacteria were pre-grown into stationary phase and spiked into Phosphate-Buffered Saline (PBS) for 24 hours, followed by transfer to Marine Broth 2216. The growth rate and cell mass were immediately recovered upon the transfer to Marine Broth 2216, suggesting that PBS serves as a valid negative control and that the bacteria remain viable and dormant in the negative control and will grow in response to nutrient availability (**fig. S4C**).

### Supplementary Note #2: Experimental and seawater sampling procedures that could impact reported cell mass measurements

Our sampling scheme and the experimental techniques that we used have the potential to alter the mass of wild and cultivated *Vibrionaceae* cells that we reported in this paper. The purpose of this note is to provide context around our sampling approach and the experimental techniques we employed, while also providing evidence we collected for how different aspects of our study could impact cell mass measurements.

**Sampling scheme**: We set out to measure the mass of individual planktonic *Vibrio* cells in seawater, which is a spatially complex environment at this scale. Bacteria cohabitate with a wide variety of organisms, particles, hydrogels, and polymeric substances. These other biological and chemical entities span a range of sizes and each can influence the aggregation of bacteria in seawater. To ensure we removed these particles, along with bacteria attached to these particles (or each other) we first used a plankton net (63µm) to remove large plankton and then fractionated biomass through a series of filters with decreasing pore size (5, 1, 0.2µm). The filters represent 3 different size fractions of biomass (5-63µm, 1-5µm, 0.2-1µm). There is no one single size cutoff where free-living bacteria solely reside, so we used a stringent cutoff of 1µm as an upper size limit to define our planktonic cell fraction. This improves our confidence that biomass passing through the previous filter is primarily made up of individual bacterial or archaeal cells. However, some larger free-living cells may not pass through the 1µm pore size filter if they contact the pore in the wrong orientation or if pores have become partially clogged with other material. Additionally, samples were flash frozen in the field, shipped back to the laboratory on dry ice, and frozen at -80C prior to fixation. While we acknowledge these sampling choices could potentially bias our results, we were still able to recover relatively rare *Vibrio* in the 0.2-1µm fraction at a reasonable relative abundance (around 1% of the total community) after sequencing the community. These sequencing results also revealed a diverse planktonic community with many ASV's (amplicon sequencing variants) classified to bacterial Families typically observed in coastal planktonic habitats (Flavobacteriaceae, SAR116, Cyanobiaceae, Actinomarinaceae, Pelagibacteriales, etc.) in the samples most representative of the full seawater community (Presort). Taken together this suggests that our sample was not severely biased by our sampling methods, but we cannot rule out that we missed some larger planktonic bacterial cells with our approach.

**Fixation of bacterial cells**: Cell fixation could alter the mass of cells, but these steps are necessary to quickly make cells metabolically inactive and preserve them from when they are removed from their environment (culture flask or seawater) until they can be measured on an SMR mass sensor. Two fixative types were used in this study, which have slightly different known impacts on cell mass. Formaldehyde is thought to work by crosslinking sulfur bonds between proteins throughout the cell, and has been previously shown to have small impacts on cell mass that range from 3-18% mass loss (*1*). Formaldehyde was used to fix pure cultures, but it has the negative feature that it is not compatible with downstream PCR amplification of DNA from formaldehyde fixed samples. Ethanol fixation is compatible with downstream PCR amplification of DNA, so it was used as the fixative for our samples that were subject to FISH, FACS, LifeScale, and sequencing (starved pure cultures of *V. cyclitrophicus* and seawater community biomass). We evaluated the impact of fixation along with other procedures in the FISH, FACS, LifeScale protocol below.

**FISH, FACS, LifeScale protocol**: Fluorescence *in situ* hybridization is a technique for labeling cells of interest with oligonucleotide probes bound to fluorophores which target ribosomal RNA. The large amount of phylogenetic information encoded in ribosomal RNA, as well as the high copy number of this molecule in cells, make it a great target for labeling cells of interest in different phylogenetic groups and reliably generating a bright signal. FISH probes are relatively small in mass when compared to bacterial cells, so are unlikely to add much mass individually. However, the number of ribosomal RNA target sequences which probes can bind in a cell are high and can vary among target cells so could theoretically impact mass. Fixation for FISH is a delicate balancing act, because one must ensure that cells remain permeable enough for oligonucleotide probes bound to fluorophores to cross the cell envelope and enter the cytoplasm. Formaldehyde fixed cells are more rigid and can become impermeable, while ethanol fixed cells have increased permeability and risk losing additional mass. We chose to use ethanol fixation for our experiments. While FACS is not likely to alter the mass of individual cells directly, there are biases that FACS sorting could introduce. For instance, we are more likely to observe brighter cells than dimmer cells with FACS and larger cells are more likely to be labeled by FISH so this could bias our sorted sample to larger cells. The physical process of sorting likely exposes cells to shear stress in the fluid streams used in flow cytometry and it is unclear if certain cells would be more likely to lyse or burst and bias their survival after sorting. We reasoned that experiments where we could compare a reference sample of living *Vibrio* cells with a known mass distribution to cells from the same sample that were frozen, filtered, fixed, FISHed, and FACS sorted would provide the best assessment of how the mass of wild *Vibrionaceae* from seawater samples would be impacted by the full protocol.

We performed experiments with *V. cyclitrophicus* 1G07 cells starved for 20 hours in PBS as our reference sample. We measured the mass of starved cells when they were alive and confirmed these mass distributions matched our previous SMR cultivation experiments with living cells inoculated and starved in PBS. We then mimicked the seawater sampling procedure in these pure cultures by filtering, freezing, and fixing filtered samples in the same way as was done in the field (excluding size fractionation, since all starved cells would pass through all of the pre-filter pore sizes used in the field to exclude eukaryotes and marine particles). We then performed FISH and FACS and measured the mass of starved cells (**fig. S10**) and found that the average mass of sorted cells significantly decreased compared to their live mass ($p$ = 0.0135). Sorted cells weighed 39.83 fg while live cells weighed 48.55fg, indicating the whole procedure led to approximately 17.96% of biomass lost. We used this mass loss percentage to correct both the starved cell mass and the wild *Vibrionaceae* cell mass in **Fig. 4B**, since the mass was being compared with other living *Vibrio* isolates growing in seawater and the growth law. While we believe this correction is the best way to compare across these datatypes, we acknowledge this may not be the choice others would make. We can say for certain that our sorted wild *Vibrionaceae* population is larger than the sorted starved *V. cyclitrophicus* 1G07 cells since the difference between them is significant when no correction is applied ($p = 0.002$).

We also observed that replicate 2 of our sorted *Vibrio* population from 2023 seawater had a lower percent of *Vibrionaceae* in our sequencing validation. Several of our sorted samples, such as the sorted *Vibrio* and sorted false positive populations, have low total biomass which can make them prone to nucleic acid contamination during 16S rRNA amplicon sequencing (*2*). We therefore sequenced sorting fluid controls and DNA extraction controls to assess for contamination in our sorting and sequencing protocols. Our sort fluid controls and kit controls contained a total of 44 ASV's. To assess if these possible contaminant sequences have impacted our sequencing results, we performed an *in silico* sequence "decontamination" by removing all 44 of these ASV's from all other samples and compared this to the same dataset without this "decontamination" step (**fig. S9A, B**). The replicate 2 sorted *Vibrio* sample had major differences in these 2 datasets, with the proportion of sequences classified as *Vibrionaceae* reaching 64.8% of all sequences in the decontaminated dataset while *Vibrionaceae* were only 34.6% of all sequences in the dataset that was not decontaminated. The other sorted *Vibrio* replicates had much smaller improvements in the total proportion of *Vibrionaceae*, and all of the replicate presort samples containing the full seawater fractionated biomass community were minimally impacted by decontamination. While this result suggests that sample 2 may have experienced nucleic acid contamination, these nucleic acids could plausibly come from either free DNA or contaminating cells that might have been present for mass measurements. We assessed the mass distributions of each replicate (**fig. S9C**) and it is apparent that replicate 2 did have fewer measured cells than the other replicates but a similar mass distribution, which aligns with this replicate having the lowest number of reads, compared to the remaining replicates. Out of an abundance of caution we compared how the average mass of wild *Vibrionaceae* would be impacted by the exclusion of rep 2 in two ways. First, we compared the mean of the median uncorrected buoyant mass for all reps (50.9 fg) to the mean of the median uncorrected buoyant mass for only reps 1 & 3 (49.5 fg) and found only a 1.4 fg difference. Second, we compared the median uncorrected buoyant mass of all replicates (50.2 fg) to the median uncorrected buoyant mass of only replicates 1 & 3 (49.6 fg) and found only a 0.6 fg difference between samples. This magnitude of difference does not impact our conclusions so therefore, we report the median mass of all replicates as our best estimate of the sorted *Vibrio* mass and acknowledge that there is a small chance that contaminant non-*Vibrio* cells could impact the mass for wild *Vibrionaceae* we report. We also acknowledge that none of our replicate sorted *Vibrio* samples was enriched to 100% classified *Vibrionaceae* sequences, so some contaminant cells could plausibly impact all our results. We thus investigated the sorted cell samples under fluorescence microscopy with DAPI to counterstain the nucleic acids, while also taking images that should capture the FISH probes specific for either *Vibrionaceae* or most Bacteria. In these representative images (**fig. S8**) we observed that sorted *Vibrio* were stained with both FISH probes and DAPI, while sorted Bacteria were not stained with the *Vibrionaceae* probes. Taken together, all of these results suggest our FISH protocol worked as expected and we did not have significant numbers of contaminant cells or nucleic acids after FACS.

**Varying fluid density among samples**: The buoyant mass of cells is proportional to the density difference between the cells and the fluid in which the cells are measured. In this study we compare samples in different seawater samples with unknown fluid densities to two types of samples with known fluid density: FACS sorted samples resuspended in sheath fluid from sorting (effectively PBS) and pure culture *Vibrio* measured in PBS. Seawater fluid density itself is not constant, but is likely to vary with salinity and other oceanographic parameters that were not measured with all samples in this study. To estimate the impact by different fluid density, we first measured the dry density of a formaldehyde fixed *Vibrio* cyclitrophicus population grown in ASW+Succinate to be 1.46 g/ml, using our previous reported method (*3*). Given the density of PBS (approximately 1.004 g/ml) and assuming the seawater density (approximately 1.026 g/ml), we then estimated that the difference between two fluids would only lead to about 5% difference in measured cell buoyant mass. Rather than attempt to convert all of our buoyant mass measurements to one common fluid density, we have decided to report buoyant mass measurements unaltered and also include the corresponding fluid densities for these measurements when they are known.

### Reference


1. B. R. K. Roller, C. Hellerschmied, Y. Wu, T. P. Miettinen, A. L. Gomez, S. R. Manalis, M. F. Polz, Single-cell mass distributions reveal simple rules for achieving steady-state growth. *mBio* **14**, e01585-23 (2023).

2. N. Fierer, P. M. Leung, R. Lappan, R. Eisenhofer, F. Ricci, S. I. Holland, N. Dragone, L. L. Blackall, X. Dong, C. Dorador, B. C. Ferrari, J. Goordial, S. P. Holmes, F. Inagaki, T. Korem, S. S. Li, T. P. Makhalanyane, J. L. Metcalf, N. Nagarajan, W. D. Orsi, E. R. Shanahan, A. W. Walker, L. S. Weyrich, J. A. Gilbert, A. D. Willis, B. J. Callahan, A. Shade, J. Parkhill, J. F. Banfield, C. Greening, Guidelines for preventing and reporting contamination in low-biomass microbiome studies. *Nat Microbiol* **10**, 1570–1580 (2025).

3. F. Feijó Delgado, N. Cermak, V. C. Hecht, S. Son, Y. Li, S. M. Knudsen, S. Olcum, J. M. Higgins, J. Chen, W. H. Grover, S. R. Manalis, Intracellular Water Exchange for Measuring the Dry Mass, Water Mass and Changes in Chemical Composition of Living Cells. *PLoS ONE* **8**, e67590 (2013).
